# Modeling the effect of daytime duration on the biosynthesis of terpenoid precursors

**DOI:** 10.1101/2024.07.15.603555

**Authors:** Oriol Basallo, Abel Lucido, Albert Sorribas, Alberto Marin-Sanguino, Ester Vilaprinyo, Emilce Martinez, Abderrahmane Eleiwa, Rui Alves

## Abstract

Terpenoids are valued chemicals in the pharmaceutical, biotechnological, cosmetic, and biomedical industries. Biosynthesis of these chemicals relies on polymerization of Isopentenyl di-phosphate (IPP) and/or dimethylallyl diphosphate (DMAPP) monomers, which plants synthesize using a cytosolic mevalonic acid (MVA) pathway and a plastidic methyleritritol-4-phosphate (MEP) pathway. Circadian regulation affects MVA and MEP pathway activity at three levels: substrate availability, gene expression of pathway enzymes, and utilization of IPP and DMAPP for synthesizing complex terpenoids. There is a gap in understanding the interplay between the circadian rhythm and the dynamics and regulation of the two pathways. In this paper we create a mathematical model of the MVA and MEP pathways in plants that incorporates the effects of circadian rhythms. We then used the model to investigate how annual and latitudinal variations in circadian rhythm affect IPP and DMAPP biosynthesis. We found that, despite significant fluctuations in daylight hours, the amplitude of oscillations in IPP and DMAPP concentrations remains stable, highlighting the robustness of the system. We also examined the impact of removing circadian regulation from different parts of the model on its dynamic behavior. We found that regulation of pathway substrate availability alone results in higher sensitivity to daylight changes, while gene expression regulation alone leads to less robust IPP/DMAPP concentration oscillations. Our results suggest that the combined circadian regulation of substrate availability, gene expression, and product utilization, along with MVA- and MEP-specific regulatory loops, create an optimal operating regime. This regime maintains pathway flux closely coupled to demand and stable across a wide range of daylight hours, balancing the dynamic behavior of the pathways and ensuring robustness in response to cellular demand for IPP/DMAPP.

## 1 Introduction

Terpenoids are a family of molecules with more than 22,000 different natural products (Harborne *et al*., 1991). Many family members have crucial biological functions. For example, in plants, they work as hormones (gibberellin, abscisic acid, etc.), photosynthetic pigments (chlorophyll, phytol, and carotenoids), electron carriers (ubiquinone, plastoquinone), mediators of the assembly of polysaccharides (polyprenyl phosphates) and structural components of membranes (phytosterols).

They are also used for other purposes, such as antibiotics, herbivore repellents, toxins and pollinator attractants (Mcgarvey and Croteau, 1995).

Plants synthesize terpenoids from two metabolic precursors: Isopentenyl di-phosphate (IPP) and dimethylallyl diphosphate (DMAPP). Two compartmentally separated pathways synthesize these precursors (Figure 1). The mevalonic acid (MVA) pathway converts acetyl-CoA (Ac-CoA) to IPP and DMAPP. This pathway is mostly cytosolic, with a couple of reactions taking place in the peroxisome. IPP and DMAPP are then used in the synthesis of phytosterols and ubiquinone (Mcgarvey and Croteau, 1995). The enzyme 3-hydroxy-3-methylglutaril-CoA reductase (HMGR) is a key enzyme in the regulation of the MVA pathway (Schaller *et al*., 1995). The methyleritritol-4- phosphate (MEP) pathway is compartmentalized in plastids and is responsible for the production of carotenoids, lateral chains of chlorophylls, plastoquinone, abscisic acid (ABA) and tocopherols (vitamin E, precursors and derivatives) (Eisenreich *et al*., 2001).

**Figure 1.**
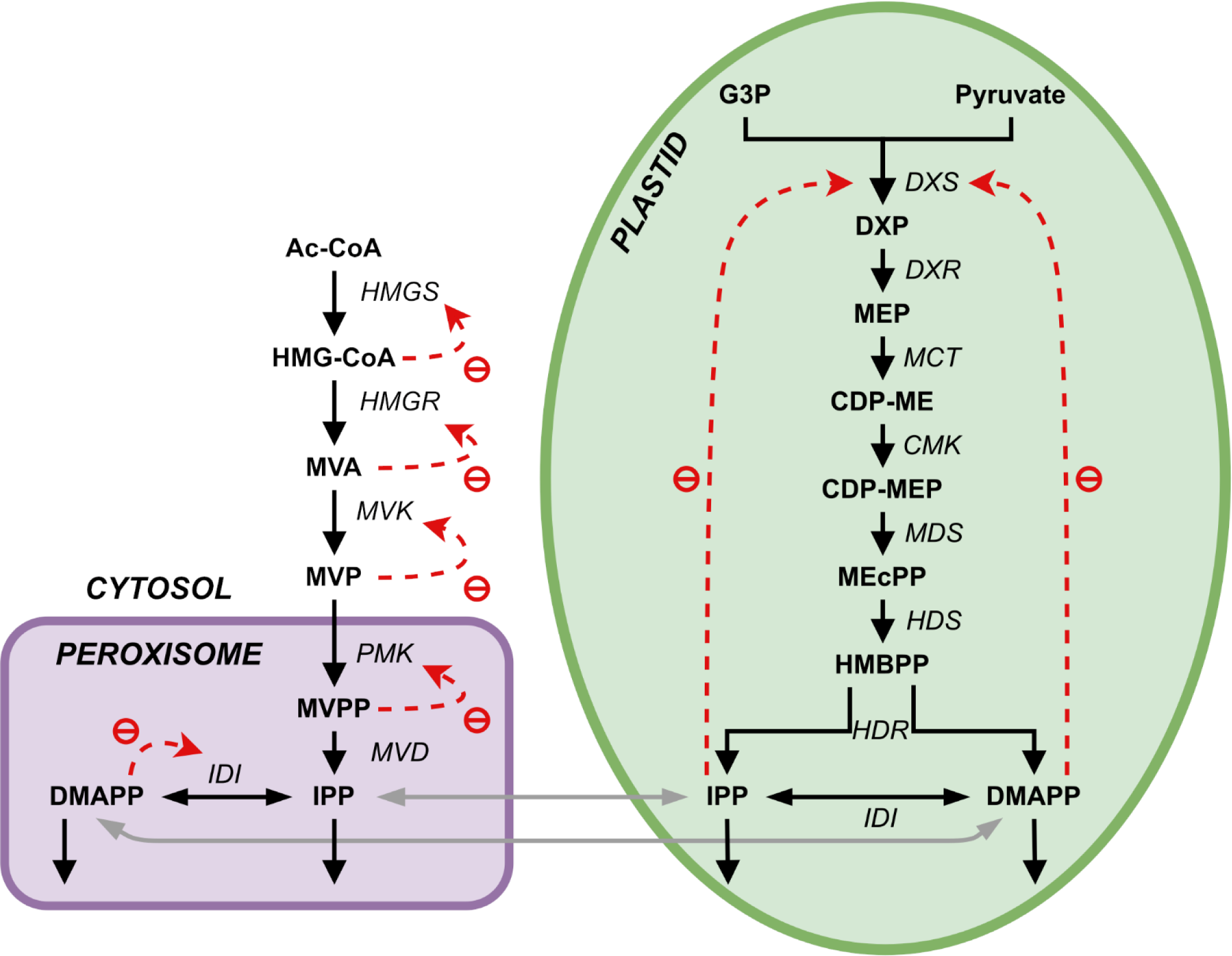
Representation of the two terpenoid biosynthesis pathways plus the ectopic pathway, the MVA pathway (left, cytosol and peroxisome) and the MEP pathway (right, plastid). DXR: DXP reductoisomerase; MCT: 2-C-methyl-D-erythrtle 4- phosphate cytidylyl transferase; CDP-ME: 4-(Citidine 5’-difosfo)-2-C-methyl-D-eritritol; CMK: 4-difosfocitidil-2-C-methyl-D-erythrtol kinase; CDP-MEP: 2-Fosfo-4-(cytidine 5’- diphospho)-2-C-methyl-D-eritritol; MDS: 2-C-methyl-D-eritritol 2,4-cyclodifosphate synthase; MEcPP: 2-C-methyl-D-eritritol 2,4-cycdiphosphate; HDS: 4-hydroxy-3-methylbut-2-en-1-il diphosphate synthase; HMBPP: 4-hydroxy-3-methylbut-2-in-1-il diphosphate; HDR: 4- hydroxy-3-methylbut-2-en-1-il diphosphate reductase; IDI: isopentenyl diphosphate Delta- isomerase; PhyPP: phytyl diphosphate.

While metabolite tracing indicates that each of the two pathways is responsible for the production of a subset of terpenoid compounds downstream, there is evidence of crosstalk between them (Hemmerlin *et al*., 2003, 2012; Hemmerlin, 2013), with some of intermediates in both pathways diffusing between the cytosol and the plastid (Bick and Lange, 2003; Laule *et al*., 2003; Hemmerlin *et al*., 2003, 2006). The first intermediate of the MEP pathway, DXP, can diffuse between the plastid and the cytoplasm (Hemmerlin *et al*., 2003; Page *et al*., 2004; Lange *et al*., 2015). At the level of IPP and DMAPP, this exchange was measured to occur mainly in the plastid-to-cytoplasm direction, promoted by a one-way symport system (Bick and Lange, 2003; Dudareva *et al*., 2005). The direction of this metabolic exchange between cellular compartments may depend on physiological state and species. There is lack of convincing evidence that other intermediates of both pathways can diffuse between the two compartments (Hemmerlin, 2013), and it would be interesting to understand what the effect of losing this exchange might have on the production of IPP/DMAPP in each compartment.

Several studies used mathematical modeling to elucidate how using synthetic biology to modify terpenoid metabolism might lead to changes in the regulatory and dynamic behavior of that metabolism. When it comes to microorganisms, for example, a kinetic model of the MVA pathway in *E. coli* using parameters from the literature correctly predicts expression and inhibition changes (Weaver *et al*., 2015). Petri nets were also used to model the integration of both pathways in yeast (Baadhe *et al*., 2012). Another example is a mathematical model of the MEP pathway in the malaria parasite *P. falciparum* (Singh and Ghosh, 2013). This ODE model was used to investigate the regulation of the pathway and to predict the effects of genetic manipulations on the production of isoprenoids with the addition of *in silico* inhibitors.

Understanding the regulation of terpenoid biosynthesis in plants is an important biological issue that has relevance also for synthetic biology (Hemmerlin *et al*., 2012; Liao *et al*., 2016; Tetali, 2019; Zhou and Pichersky, 2020; Mcgarvey and Croteau, 1995; Harborne *et al*., 1991). One of the master regulators of metabolism in plants is the circadian rhythm imposed by earth’s rotation. Mathematical models were also used to study the effect of that rhythm on the dynamics of terpenoid precursor biosynthesis in the MEP pathway of peppermint leaves (Rios-Estepa *et al*., 2010) and *Arabidopsis* (Pokhilko *et al*., 2015) (Neiburga *et al*., 2023). These three models focus on and include metabolites downstream of the precursors IPP and DMAPP, and do not include the intermediates upstream.

Measurements of volatile terpenoid emissions in plants show circadian oscillations with peaks during the day and a general decrease at night (Loivamäki *et al*., 2007; Zheng *et al*., 2017; Zeng *et al*., 2017; Mu *et al*., 2022; Picazo-Aragonés *et al*., 2020). These oscillations are driven by light regulation and internal circadian clocks in the MEP pathway. There is evidence of light regulated DXR expression and internal circadian clock regulation downstream via regulation of isoprene synthase. However, the internal clock oscillations are best maintained when coordinated by light cycles (Loivamäki *et al*., 2007). Gene expression analysis has shown that MEP pathway genes *DXS*, *DXR*, *CMK*, *MCT*, *MDS*, *HDS*, *HDR* and *IDI* present circadian oscillation, as well as many other downstream genes involved in carotene, tocopherol and other phytohormone biosynthesis, their expression levels rising during the day and decreasing through the night (Zheng *et al*., 2017; Covington *et al*., 2008).

The key genes for circadian regulation that are conserved across many plant species are *CCA1*, *LHY*, *TOC1* and *ZTL* (McClung, 2013; Nagel and Kay, 2012). *CCA1* and *LHY* inhibit *TOC1* and vice versa, while *ZTL* tags *TOC1* for degradation under blue light, which favors the morning components *CCA1* and *LHY*. These morning components then upregulate MEP pathway genes (Picazo-Aragonés *et al*., 2020; Alabadí *et al*., 2001; Yon *et al*., 2017).

Currently, the use of mathematical models to explore the dynamics and regulatory mechanisms within the MVA pathway, and more critically, the intricate interplay between the MEP and MVA pathways, remains in its embryonic stages (Basallo *et al*., 2023). To our knowledge, (Basallo *et al*., 2023) provide the only example where this is done, creating and analyzing a comprehensive model that delineates these pathways in plants, accounting for all intermediates leading to the crucial precursors IPP and DMAPP. This lack of attention may stem from the predominant focus on the MEP pathway for synthesizing chemical species of interest in plants, while MVA-derived products, equally significant, have been predominantly studied in microorganisms. As such, integrating both pathways into mathematical models that can be used as tools to study the integrated dynamics and regulation of both pathways stands as a paramount necessity for comprehensive understanding and exploration in plant biochemistry and metabolism.

In this paper, we adapted the mathematical model of the MVA and MEP pathways in *Oryza sativa* (rice) (Basallo *et al*., 2023) to account for the regulation of the kinetic activity in those pathways by the circadian rhythm. We then used the model to investigate how the changes in that rhythm over the year and at different latitudes affect IPP and DMAPP biosynthesis. Finally, we performed a set of *in silico* experiments where we removed circadian regulation from different parts of the model to investigate how the various regulatory loops contribute to the dynamic behavior of IPP and DMAPP biosynthesis. This enabled us to examine how different regulatory designs for the network (the network genotype) influence the adaptation of the dynamic behavior of IPP/DMAPP biosynthesis (the network phenotype) to the changes in daylight hours that occur due to the Earth’s circadian rhythm.

## 2 Materials and methods

### 2.1 Mathematical modeling formalism

To model the biosynthesis of IPP/DMAPP, we employed systems of ODEs. The saturating formalism was employed as the mathematical framework to depict the flux dynamics (Sorribas *et al*., 2007; Alves *et al*., 2008). This formalism approximates the kinetics of any given reaction to parameters that have biochemical interpretations in enzyme kinetics. In this formalism, we approximate the rate of a reaction in an inverse space at an operating point by:

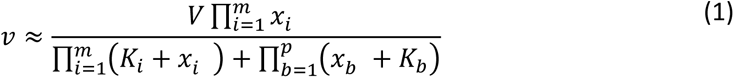

V parameters represent apparent saturation rate constants for the reactions. Ki parameters represent apparent binding constants for the substrate(s) or inhibitor(s) of the reaction. While no activators were considered in our model, these can also be included using this formalism.

### 2.2 Mathematical models for the MVA and MEP pathways

We adapted the model published in (Basallo *et al*., 2023), and summarized in Tables 1–3. There, the pathways are modelled using the canonical reaction set for each pathway (shown in Figure 1), extracted from KEGG and the literature consensus. Exchange of IPP and DMAPP between the cytosol (produced by the MVA pathway) and the plastid (produced by the MEP pathway) is modelled by considering that, under physiological conditions, IPP and DMAPP mostly flow from the plastid into the cytosol (Bick and Lange, 2003). This is implemented by setting the flow from the plastid to the cytosol to be ten times the rate of the import to the cytosol. Bick and Lange (Bick and Lange, 2003) also reported that other pathway intermediates were not actively transported between the two compartments. Table 3 summarizes all reactions of material interchanged between plastid and cytosol.

**Table 1.** MVA pathway reactions that were considered in the model.

| MVA pathway (cytoplasm) | Rate Expression | Rate |
| --- | --- | --- |
| Acetoacetyl-CoA<sub>cyt</sub> → HMG-CoA<sub>cyt</sub> | $(V_{max1} HMGS \text{ acetoacetyl-CoA}_{cyt}) / (\text{acetoacetyl-CoA}_{cyt} + Km1 (1 + (HMG-CoA_{cyt})/Ki1) )$ | $r_1$ |
| HMG-CoA<sub>cyt</sub> → MVA<sub>cyt</sub> | $\frac{V_{max2} HMGR HMG-CoA_{cyt}}{HMG-CoA_{cyt} + Km2 \left( 1 + \frac{MVA_{cyt}}{Ki2} \right)}$ | $r_2$ |
| MVA<sub>cyt</sub> → MVP<sub>cyt</sub> | $\frac{V_{max3} MVK MVA_{cyt}}{MVA_{cyt} + Km3 \left( 1 + \frac{MVP_{cyt}}{Ki31} + \frac{FPP_{cyt}}{Ki32} + \frac{GPP_{cyt}}{Ki33} + \frac{GGPP_{cyt}}{Ki34} + \frac{PhyPP_{cyt}}{Ki35} \right)}$ | $r_3$ |
| MVP<sub>cyt</sub> → MVPP<sub>cyt</sub> | $\frac{V_{max4} PMK MVP_{cyt}}{MVP_{cyt} + Km4 \left( 1 + \frac{MVPP_{cyt}}{Ki4} \right)}$ | $r_4$ |
| MVPP<sub>cyt</sub> → IPP<sub>cyt</sub> | $\frac{V_{max5} MVD MVPP_{cyt}}{MVPP_{cyt} + Km5}$ | $r_5$ |
| IPP<sub>cyt</sub> → DMAPP<sub>cyt</sub> | $\frac{V_{max6} IDI IPP_{cyt}}{IPP_{cyt} + Km6 \left( 1 + \frac{DMAPP_{cyt}}{Ki6} \right)}$ | $r_6$ |
| DMAPP<sub>cyt</sub> → IPP<sub>cyt</sub> | $\frac{V_{max7} IDI DMAPP_{cyt}}{DMAPP_{cyt} + Km7}$ | $r_7$ |
| 2 IPP<sub>cyt</sub> + 4 DMAPP<sub>cyt</sub> → | $k1 IPP_{cyt} DMAPP_{cyt}$ | $r_{20}$ |

**Table 2.**
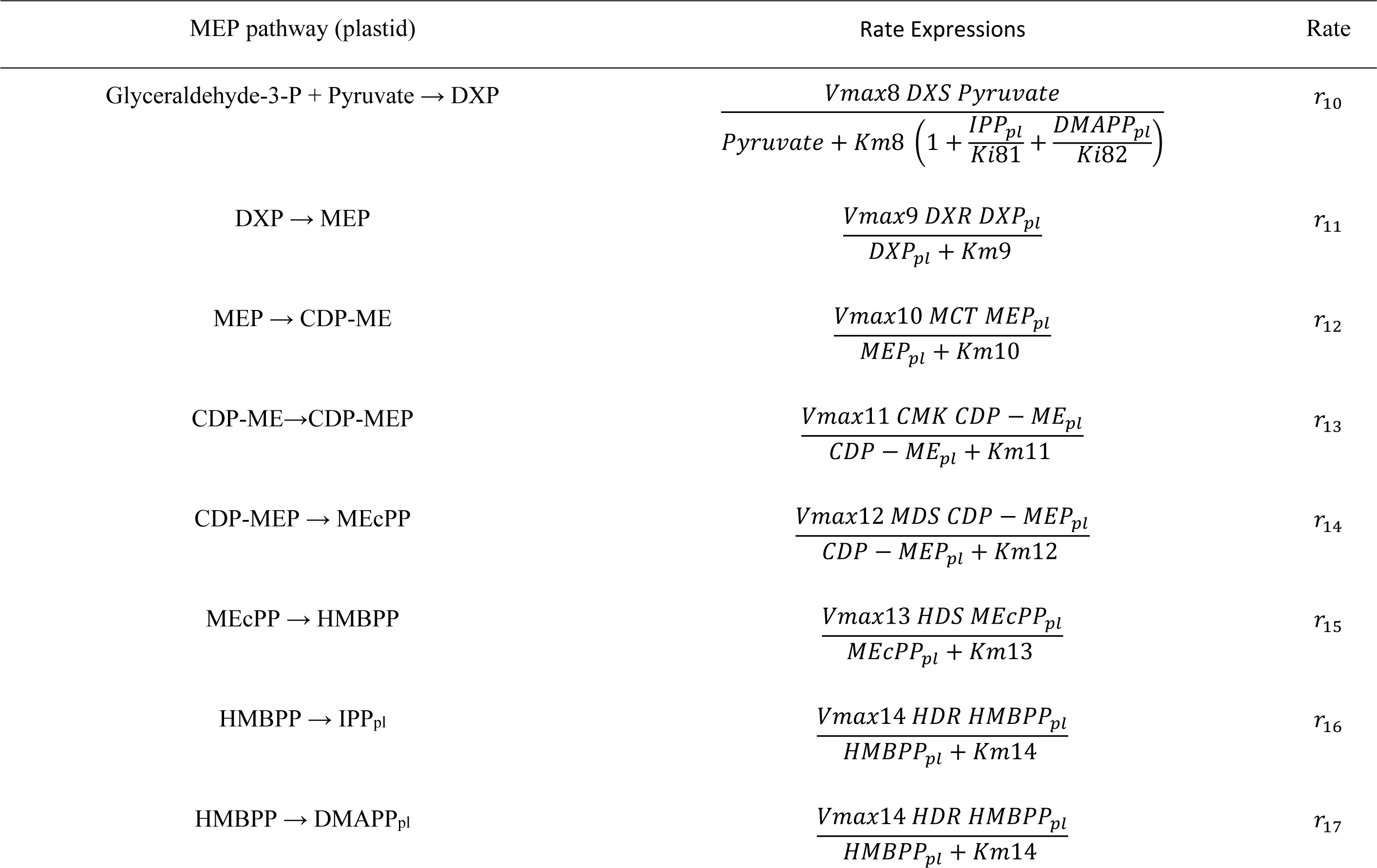
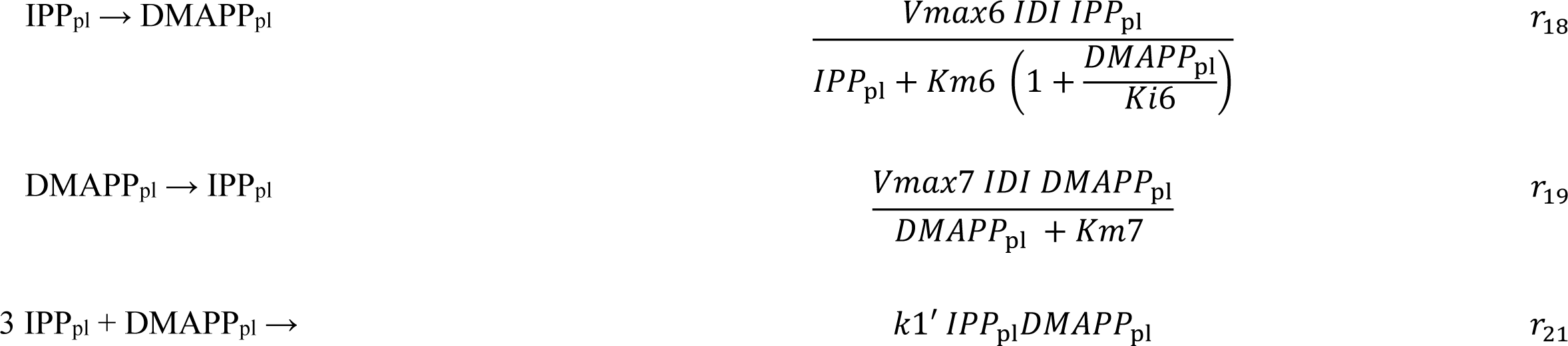
MEP pathway reactions that were considered in the model.

**Table 3.** Exchange of MVA and MEP intermediates between the cytosol and the plastid.

| Exchange of material between cytoplasm and plastid | Rate expressions | Rate |
| --- | --- | --- |
| $IPP_{\text{cyt}} \rightarrow IPP_{\text{pl}}$ | $k2 IPP_{\text{cyt}}$ | $r_8$ |
| $IPP_{\text{pl}} \rightarrow IPP_{\text{cyt}}$ | $k3 IPP_{\text{pl}}$ | $r_9$ |
| $DMAPP_{\text{cyt}} \rightarrow DMAPP_{\text{pl}}$ | $k2' DMAPP_{\text{cyt}}$ | $r_{22}$ |
| $DMAPP_{\text{pl}} \rightarrow DMAPP_{\text{cyt}}$ | $k3' DMAPP_{\text{pl}}$ | $r_{23}$ |

We modelled the kinetics of each step, as well as those for the exchange fluxes of IPP and DMAPP between cytoplasm and plastid, using the rate expressions in Tables 1-3. We assume that the organism maintains homeostasis of Acetyl-CoA and Acetoacetyl-CoA. Table 4 presents the kinetic constants for each reaction, retrieved from (Basallo *et al*., 2023) and from the primary literature. Table 5 collects the concentrations for the independent variables. Hereafter, we refer to this model as Model A.

**Table 4.**
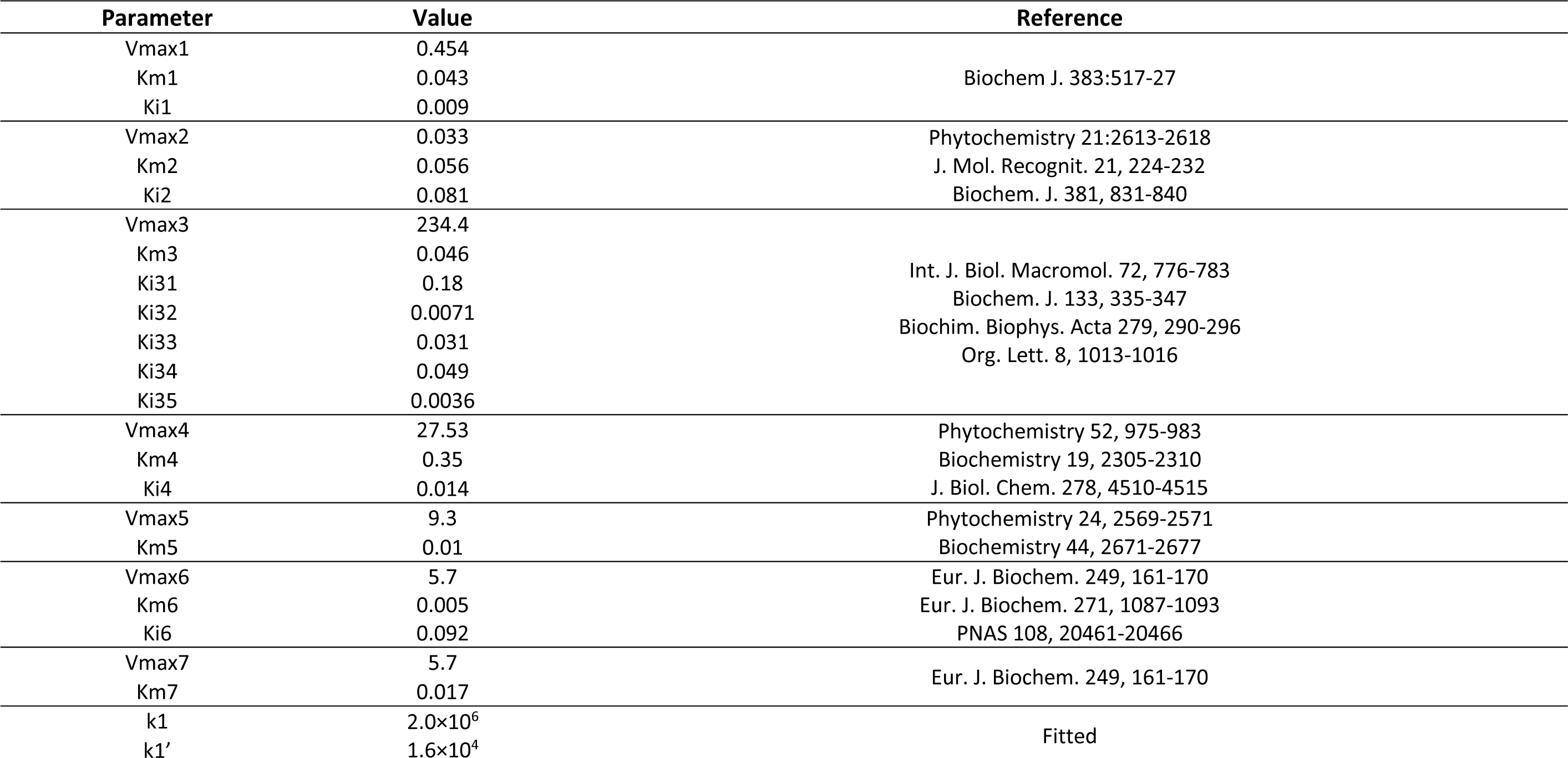
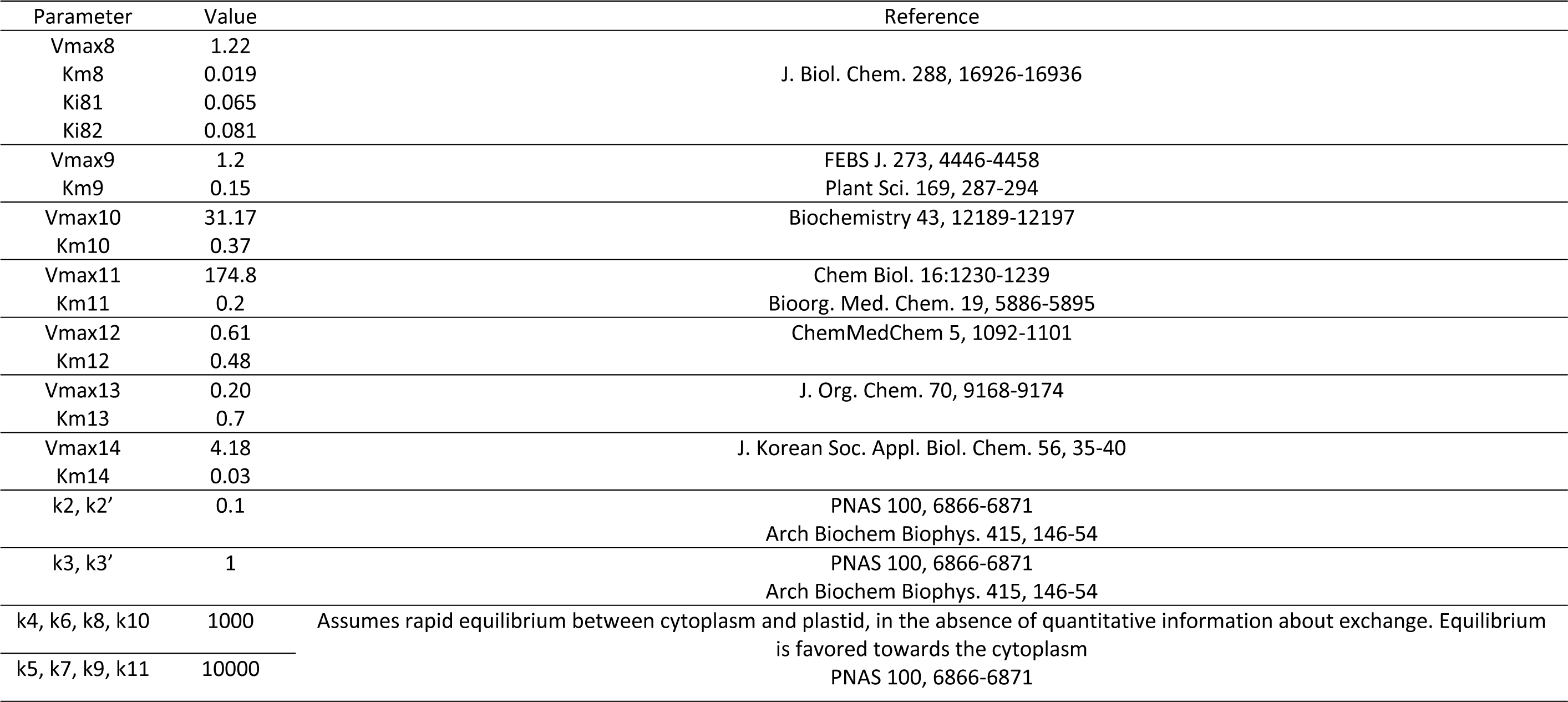
Kinetic Parameters. All concentration units in mM. All time units in s^-1^.

| Parameter | Value | Reference |
| --- | --- | --- |
| Vmax1 | 0.454 | Biochem J. 383:517-27 |
| Km1 | 0.043 |  |
| Ki1 | 0.009 |  |
| Vmax2 | 0.033 | Phytochemistry 21:2613-2618 |
| Km2 | 0.056 | J. Mol. Recognit. 21, 224-232 |
| Ki2 | 0.081 | Biochem. J. 381, 831-840 |
| Vmax3 | 234.4 | Int. J. Biol. Macromol. 72, 776-783<br>Biochem. J. 133, 335-347<br>Biochim. Biophys. Acta 279, 290-296<br>Org. Lett. 8, 1013-1016 |
| Km3 | 0.046 |  |
| Ki31 | 0.18 |  |
| Ki32 | 0.0071 |  |
| Ki33 | 0.031 |  |
| Ki34 | 0.049 |  |
| Ki35 | 0.0036 |  |
| Vmax4 | 27.53 | Phytochemistry 52, 975-983 |
| Km4 | 0.35 | Biochemistry 19, 2305-2310 |
| Ki4 | 0.014 | J. Biol. Chem. 278, 4510-4515 |
| Vmax5 | 9.3 | Phytochemistry 24, 2569-2571 |
| Km5 | 0.01 | Biochemistry 44, 2671-2677 |
| Vmax6 | 5.7 | Eur. J. Biochem. 249, 161-170 |
| Km6 | 0.005 | Eur. J. Biochem. 271, 1087-1093 |
| Ki6 | 0.092 | PNAS 108, 20461-20466 |
| Vmax7 | 5.7 | Eur. J. Biochem. 249, 161-170 |
| Km7 | 0.017 |  |
| k1 | 2.0×10<sup>6</sup> | Fitted |
| k1' | 1.6×10<sup>4</sup> |  |

Table 4 (continued) Kinetic Parameters. All concentration units in mM. All time units in s<sup>-1</sup>.
| Parameter | Value | Reference |
| --- | --- | --- |
| Vmax8 | 1.22 | J. Biol. Chem. 288, 16926-16936 |
| Km8 | 0.019 |  |
| Ki81 | 0.065 |  |
| Ki82 | 0.081 |  |
| Vmax9 | 1.2 | FEBS J. 273, 4446-4458 |
| Km9 | 0.15 | Plant Sci. 169, 287-294 |
| Vmax10 | 31.17 | Biochemistry 43, 12189-12197 |
| Km10 | 0.37 |  |
| Vmax11 | 174.8 | Chem Biol. 16:1230-1239 |
| Km11 | 0.2 | Bioorg. Med. Chem. 19, 5886-5895 |
| Vmax12 | 0.61 | ChemMedChem 5, 1092-1101 |
| Km12 | 0.48 |  |
| Vmax13 | 0.20 | J. Org. Chem. 70, 9168-9174 |
| Km13 | 0.7 |  |
| Vmax14 | 4.18 | J. Korean Soc. Appl. Biol. Chem. 56, 35-40 |
| Km14 | 0.03 |  |
| k2, k2' | 0.1 | PNAS 100, 6866-6871 |
|  |  | Arch Biochem Biophys. 415, 146-54 |
| k3, k3' | 1 | PNAS 100, 6866-6871 |
|  |  | Arch Biochem Biophys. 415, 146-54 |
| k4, k6, k8, k10 | 1000 | Assumes rapid equilibrium between cytoplasm and plastid, in the absence of quantitative information about exchange. Equilibrium is favored towards the cytoplasm |
| k5, k7, k9, k11 | 10000 |  |
|  |  | PNAS 100, 6866-6871 |

**Table 5.** Concentration of independent Variables (Albe *et al*., 1990).

| *Metabolite* | *Concentration (mM)* |
| --- | --- |
| *Ac-CoA* | 0.350 |
| *G3P* | 0.006 |
| *Pyruvate* | 1.600 |

### 2.3 Stability analysis

We assess stability of the steady states by calculating the eigenvalues of the Jacobian matrix of the ODE system, which are complex numbers (Voit, 2013). If the real parts of all eigenvalues are negative, the system is stable. Otherwise, the system is unstable. The Jacobian matrix is constructed by taking the partially derivatives of the right-hand side of the ODEs (*f_i_*) with respect to each state variable (x_*j*_), as shown in Eq. 2.

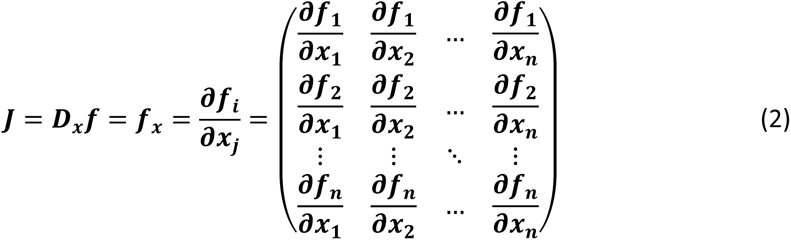

### 2.4 Sensitivity analysis

Logarithmic sensitivity analysis of the system was performed by calculated logarithmic, or relative, steady-state parameter sensitivities, which measure the “relative change in a system variable (X) that is caused by a relative change in a parameter (p)” (Voit, 1991):

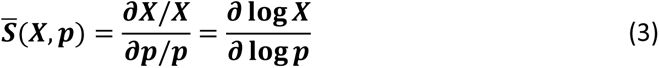

We also calculated the steady state sensitivities of the aggregate input flux and aggregate output flux of each metabolite to the model parameters using Eq (4).

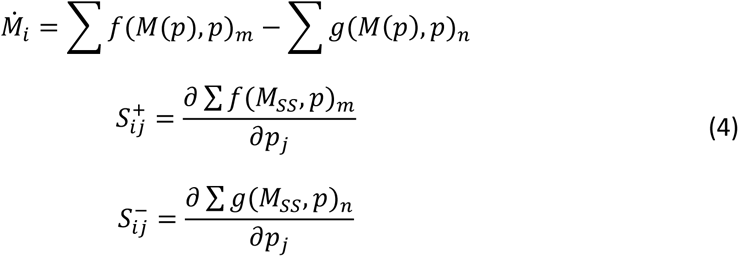

### 2.5 Modeling the circadian rhythm

To model the effect of the circadian oscillation on the dynamics of Model A we adapted the approach used by (Pokhilko *et al*., 2014, 2015), described by Eq 5:

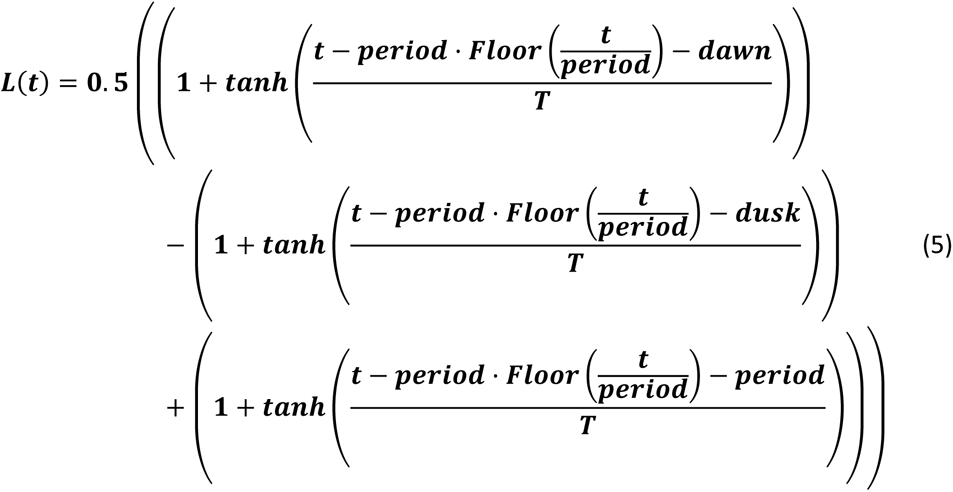

***L***(***t***) can have values between 0 (no light) and 1 (maximal diurnal light intensity). The *Floor* function returns the greatest integer less than or equal to the input value. We set the *period* of oscillation to 24 h, the duration of the day. To model the positive effect of the presence of light on a variable *X_i_* or a parameter *p_j_* we use Eq 6:

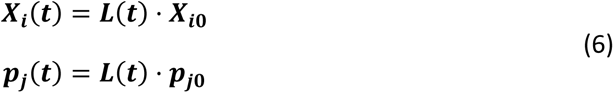

In contrast, to model the negative effect of the presence of light on a variable *X_i_*or a parameter *p_j_* we use Eq 7:

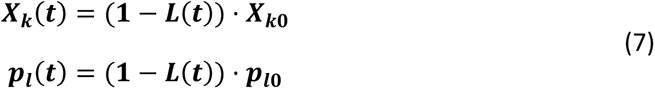

### 2.6 Modeling the effect of latitude and seasonality on the number of hours of light per day

Parameters *dawn* and *dusk* control the time of sunrise and sunset respectively. By setting *dawn* to 0 h, we can control daytime length through changing only the *dusk* parameter. The default value we use for *dusk* is 12 h. This leads to ***L***(***t***) having 12h of light and 12h of dark, which is the situation close to the equator. Increasing dusk above 12h makes ***L***(***t***) have more than 12h of daylight within the 24h period of the oscillation. Decreasing dusk below 12h makes ***L***(***t***) have less than 12h of daylight within the 24h period of the oscillation. In this way, *dusk* is a proxy for the effect of geographic latitude on the number of daylight hours within a circadian oscillation.

Parameter *T* emulates twilight duration. At *T* close to 0 h, the function approximates a step function (square wave) while progressively higher values of T lead to a less steep transition between darkness and full light levels (sinusoidal wave).

The number of daylight hours in a day changes with the seasons and with latitude. In the peak of the northern hemisphere winter, there are 0 h of daylight in the north pole and 24 h of daylight in the south pole. Supplementary Figure S1 illustrates how the L function can model the number of full daylight, twilight and night hours at different latitudes and during the duration of a single year.

### 2.7 Modeling the effects of circadian oscillations on the dynamics of the MVA and MEP pathways

There are three regulatory circadian modules we consider in our model: regulation of substrate production for the MVA end MEP pathways, regulation of gene expression in the pathways, and regulation of IPP/DMAPP consumption after they are synthesized.

At the level of substrate production, (Cockburn and McAulay, 1977) show that triose intermediates of glycolysis remain roughly constant over the circadian light cycle. In contrast, they also show that pyruvate in plants leaves can oscillate over the daylight cycle, changing over two-fold with respect to its average value. (Pokhilko *et al*., 2014, 2015) assume that the circadian light cycle controls the availability of the triose G3P, which is a precursor of the MEP pathway. Similarly, the availability of Acetyl-CoA in the cytoplasm of plant leaves also changes over the circadian light cycle (Lee *et al*., 2021; Buchanan *et al*., 2002; Taiz *et al*., 2014). In the morning, acetyl-CoA levels rise, as photosynthesis becomes more active, reaching peak levels at midday. Acetyl-CoA availability decreases during the afternoon, remaining low as photosynthesis is absent and metabolic activity is reduced during the night. We follow the experimental evidence by Cockburn and McAuly (1977), combining it with the approach used by (Pokhilko *et al*., 2014, 2015) and model the effect of circadian rhythms on the availability of MVA and MEP precursors using Eq. 8:

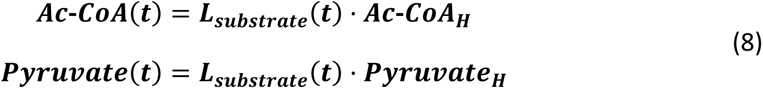

Here, ***Pyruvate _H_*** and ***Ac-CoA _H_*** represent the steady state values for pyruvate and acetyl-coA in model A. (Table 5).

Model A modified through the addition of Eq. 8 will be referred to hereafter as Model B. Model B reverts to Model A when ***L(t)*** *= 1*.

At the level of gene expression regulation, experimental evidence indicates that there is anti-phasic regulation of MVA and MEP pathway genes by daylight (Atamian and Harmer, 2016; Jin *et al*., 2021; Covington *et al*., 2008; Vranová *et al*., 2013). The data suggests that daylight activates MEP pathway genes (Jin *et al*., 2021) and deactivates MVA genes. To model this effect, we made protein activity time dependent and directly proportional to light level, using Eq 9:

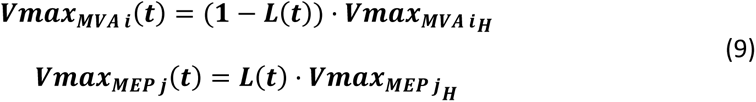

Index *i* represents each enzyme of the MVA pathway, while index *j* represents each enzyme of the MEP pathway.***Vmax_MVAiH_*** represents the basal value of ***Vmax_MVAiH_*** in Model A (Table 5).

Hereafter we will refer to the Model A modified with Eq 9 as Model C.

At the level of IPP/DMAPP consumption, and following (Pokhilko *et al*., 2014, 2015) and the experimental evidence (Covington *et al*., 2008; Loivamäki *et al*., 2007; Zheng *et al*., 2017; Zeng *et al*., 2017; Picazo-Aragonés *et al*., 2020; Mu *et al*., 2022),we model usage of IPP and DMAPP downstream of the MVA as being dependent of the circadian rhythm. In fact, the expression of genes from pathways that synthesize more complex terpenoids was observed to be coordinated to that of the genes from the MEP pathway (Jin *et al*., 2021). It is well established experimentally that plant emission of terpenoid oils, which are derived from IPP and DMAPP precursors is high during the light hours of the day and strongly decreases during nighttime. We model this effect by modifying the rate constants of reactions r20 and r21 according to Eq 10:

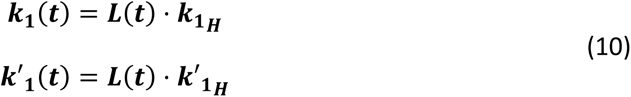

Here, ***k_1 H_*** and ***k’_1 H_*** are the values of k1 and k’1 used in Model A. Model A modified with Eq 10 will be referred to hereafter as Model D.

By combining Models B, C, and D we investigated how changes in the regulatory action of circadian rhythms might affect the dynamic behavior of the MEP and MVA pathways. This is the mathematical equivalent of mutating the genome to eliminate, create, or modify regulatory loops with the purpose of studying their effect. Thus, in Model BC we eliminate regulation of IPP/DMAPP consumption by the circadian rhythm, in Model BD we eliminate regulation of MVA and MEP gene expression by circadian rhythms, and in Model CD we eliminate regulation of pathway substrate production by circadian rhythms. Finally, we assemble the modifications of Models B, C, and D in a single model, which we refer to as Model E.

### 2.8 Model Implementation

The Mathematica code for the implementation of all models and Figures is provided in Supplementary Data File S1.

## 3 Results

### 3.1 Quality analysis of the basal model for the MEP and MVA pathways

Eq. 11 represents the joint basal mathematical model (Model A hereafter) of the MEP and MVA pathways adapted from (Basallo *et al*., 2023):

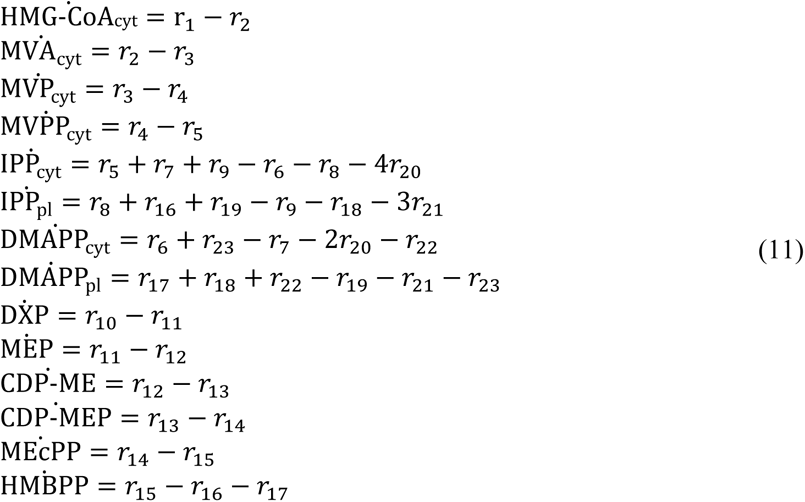

Details about the mathematical form and parameter values of each flux function r_i_ are given in Tables 1-3. Here we investigate whether the model has a positive, biological reasonable, steady state that is stable and robust. Table 6 provides the steady state concentration of the variables in the Model. The concentrations are within normal metabolite concentration ranges. In addition, the steady state is stable, having negative real parts for all the eigenvalues of the system’s Jacobean matrix (Table 7).

**Table 6.** Predicted concentrations of metabolites (mM).

| Metabolites | Model | Km range* |
| --- | --- | --- |
| HMGCoA<sub>cyt</sub> | 0.983 | [0.007 – 3.7] |
| MVA<sub>cyt</sub> | 3.5x10<sup>-5</sup> | [0.012 – 0.14] |
| MVP<sub>cyt</sub> | 3.98x10<sup>-4</sup> | [0.004 – 2] |
| MVPP<sub>cyt</sub> | 3.36x10<sup>-5</sup> | [0.001 – 0.02] |
| IPP<sub>cyt</sub> | 0.109 | [0.0057 – 0.5] |
| IPP<sub>pla</sub> | 0.0801 | [0.0057 – 0.5] |
| DMAPP<sub>cyt</sub> | 0.136 | 0.017 |
| DMAPP<sub>pla</sub> | 0.124 | 0.017 |
| DXP | 0.0133 | [0.0031 – 0.12] |
| MEP | 1.15x10<sup>-3</sup> | [0.003 – 3.26] |
| CDPME | 1.11x10<sup>-4</sup> | [0.001 – 0.2] |
| CDPMEP | 0.0920 | – |
| MECPP | 0.657 | – |
| HMBPP | 3.52x10<sup>-4</sup> | [0.006 – 0.59] |

\* Values taken from BRENDA (BRENDA Enzyme Database, 2024). The Kms refer to the reaction where the metabolite is a substrate. Kms provide a reasonable, but not definitive estimation of the physiological value for the substrate of each reaction (Savageau, 1976).

**Table 7.** Eigenvalues for the Steady State.

|  | **Real** | **Im** |
| --- | --- | --- |
| Eigenvalue1 | -928.693 | 0 |
| Eigenvalue2 | -891.953 | 0 |
| Eigenvalue3 | -872.862 | 0 |
| Eigenvalue4 | -272.249 | 0 |
| Eigenvalue5 | -83.721 | 0 |
| Eigenvalue6 | -78.084 | 0 |
| Eigenvalue7 | -16.974 | 0 |
| Eigenvalue8 | -11.845 | 0 |
| Eigenvalue9 | -6.668 | 0 |
| Eigenvalue10 | -2.011 | 0 |
| Eigenvalue11 | -0.913 | 0 |
| Eigenvalue12 | -0.545 | 0 |
| Eigenvalue13 | -0.110 | 0 |
| Eigenvalue14 | -0.031 | 0 |

The model also predicts that the concentrations of pathway intermediates are lower than those of its substrates (HMG-CoA and DXP) and end-products (DMAPP and IPP), which is another hallmark of a well-behaved biosynthetic pathway (Alves and Savageau, 2000).

Sensitivity analysis identifies the parameters to which the variables of the model are most sensitive, as described in (Sorribas *et al*., 2007; Alves *et al*., 2008). A high sensitivity of a variable to a parameter indicates that small changes in the value of that parameter might lead to large changes in the value of the variable.

To understand how sensitive the stability of the steady state is to perturbations in the parameters of the models, we calculated the logarithmic sensitivity of the steady state Jacobian eigenvalues to each parameter of the model (Table 8). The model has over eighty parameters and eigenvalues have sensitivities that are above one (in absolute values) to thirty of them. The parameters to which more eigenvalues are sensitive concentrate in reactions r2 (HMG-CoA_cyt_ → MVA_cyt_), r3 (MVA_cyt_ → MVP_cyt_), r4 (MVP_cyt_ → MVPP_cyt_), and r6 (IPP_cyt_ → DMAPP_cyt_) of the MVA pathway and reactions r10 (Glyceraldehyde-3-P + Pyruvate → DXP) and r18 (IPP_pl_ → DMAPP_pl_) of the MEP pathways.

**Table 8.**
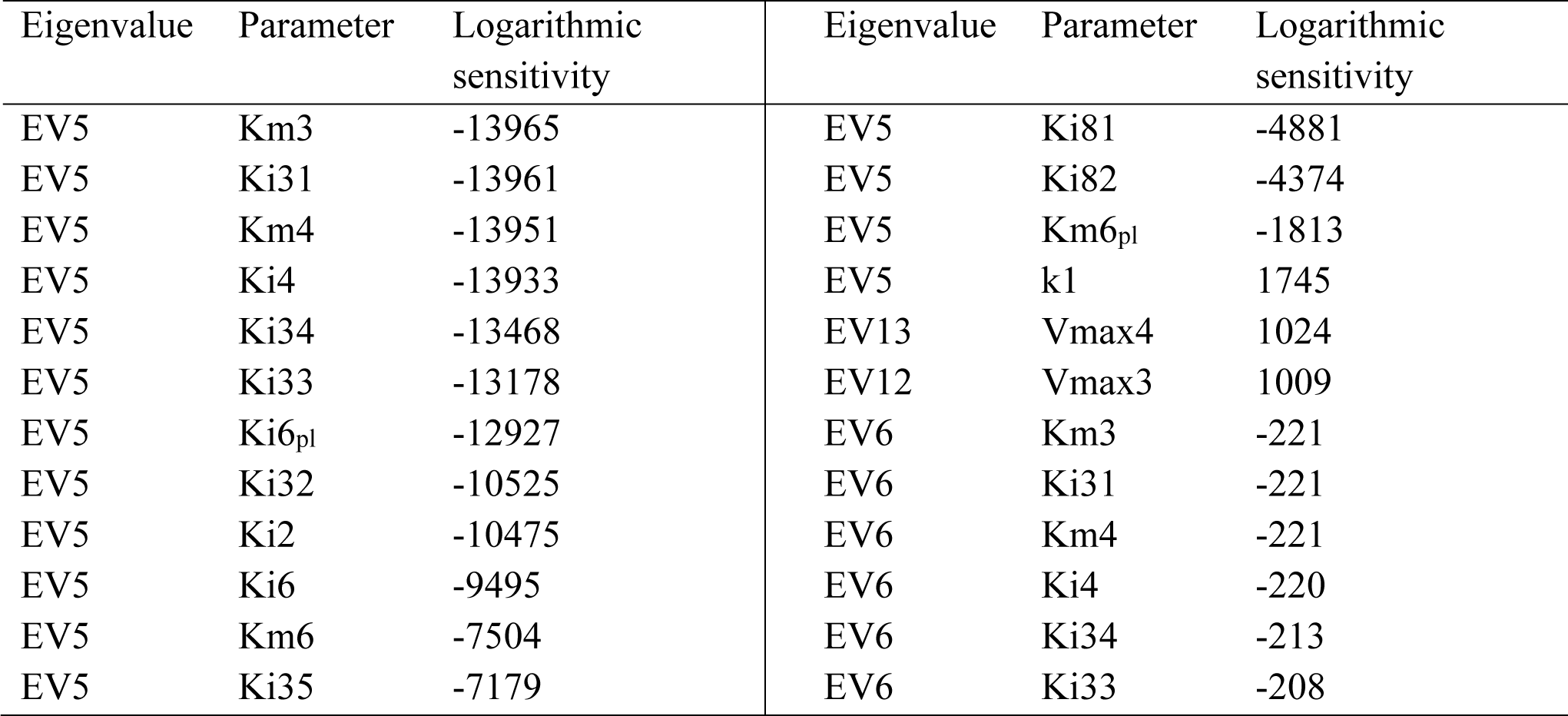
Logarithmic sensitivities of the eigenvalues to reaction parameters. Only the top 24 sensitivities with largest absolute value are shown. Full table in Supplementary Data S1.

The eigenvalues whose real part is closer to zero are most sensible to Vmax3, Vmax4, Vmax5, Km3, Ki31, and Km4, parameters from r3 (MVA_cyt_ → MVP_cyt_), r4 (MVP_cyt_ → MVPP_cyt_) and r5 (MVPP_cyt_→ IPP_cyt_).

Plausible models of biological systems have low sensitivities to most parameters (Savageau, 1976; Kitano, 2007). The logarithmic sensitivity analysis of the dependent concentrations with respect to each parameter of the model we performed shows that our model fits this quality criterion. Only 51 out of 728 sensitivities are larger than 0.5 and none is larger than 3 in absolute value (Table 9).

**Table 9.** Logarithmic sensitivities of the concentrations to reaction parameters. Only sensitivities with an absolute value larger than 0.5 are shown.

| Variable | Parameter | Sensitivity | Variable | Parameter | Sensitivity |
| --- | --- | --- | --- | --- | --- |
| HMGC<sub>CoA<sub>cyt</sub></sub> | Ki1 | 0.945 | MEP | Km8 | -0.922 |
| HMGC<sub>CoA<sub>cyt</sub></sub> | Km1 | -0.953 | MEP | Vmax8 | 1.00 |
| HMGC<sub>CoA<sub>cyt</sub></sub> | Vmax1 | 1.02 | MEP | Km10 | 1.00 |
| HMGC<sub>CoA<sub>cyt</sub></sub> | Vmax2 | -1.02 | MEP | Vmax10 | -1.00 |
| MVA<sub>cyt</sub> | Vmax2 | 0.945 | CDPME | Km8 | -0.920 |
| MVA<sub>cyt</sub> | Km3 | 1.00 | CDPME | Vmax8 | 0.999 |
| MVA<sub>cyt</sub> | Vmax3 | -1.00 | CDPME | Km11 | 1.00 |
| MVP<sub>cyt</sub> | Vmax2 | 0.948 | CDPME | Vmax11 | -1.00 |
| MVP<sub>cyt</sub> | Km4 | 1.00 | CDPMEP | Km8 | -1.09 |
| MVP<sub>cyt</sub> | Vmax4 | -1.00 | CDPMEP | Vmax8 | 1.19 |
| MVPP<sub>cyt</sub> | Vmax2 | 0.948 | CDPMEP | Km12 | 1.00 |
| MVPP<sub>cyt</sub> | Km5 | 1.00 | CDPMEP | Vmax12 | -1.19 |
| MVPP<sub>cyt</sub> | Vmax5 | -1.00 | MECPP | Ki81 | 0.584 |
| IPP<sub>cyt</sub> | Vmax6 | -0.929 | MECPP | Ki82 | 0.725 |
| IPP<sub>cyt</sub> | Vmax7 | 1.79 | MECPP | Km8 | -1.79 |
| IPP<sub>pla</sub> | Vmax6<sub>pl</sub> | -2.86 | MECPP | Vmax8 | 1.94 |
| IPP<sub>pla</sub> | Vmax7<sub>pl</sub> | 2.89 | MECPP | Km13 | 1.00 |
| IPP<sub>pla</sub> | k1' | -0.621 | MECPP | Vmax13 | -1.94 |
| DMAPP<sub>cyt</sub> | Vmax6 | 0.929 | MECPP | k1' | 0.638 |
| DMAPP<sub>cyt</sub> | Vmax7 | -1.79 | HMBPP | Km8 | -0.930 |
| DMAPP<sub>cyt</sub> | k1' | -0.500 | HMBPP | Vmax8 | 1.01 |
| DMAPP<sub>pla</sub> | Vmax6<sub>pl</sub> | 2.86 | HMBPP | Vmax14 | -0.505 |
| DMAPP<sub>pla</sub> | Vmax7<sub>pl</sub> | -2.89 | HMBPP | Km14 | 0.501 |
| DXP | Km8 | -1.00 | HMBPP | Vmax14' | -0.507 |
| DXP | Vmax8 | 1.09 |  |  |  |
| DXP | Km9 | 1.00 |  |  |  |
| DXP | Vmax9 | -1.09 |  |  |  |

DMAPP and IPP are the metabolites with the highest sensitivities. The parameters responsible for these high sensitivities are the maximum velocities of isomerization between DMAPP and IPP. In general, the parameters causing the highest sensitivities for each metabolite correspond to a reaction directly involved in producing or consuming that metabolite. In addition, the metabolites on the MEP pathway seem to share a high sensitivity to parameters from rate *r10*. DXS catalyzes this reaction, where G3P and pyruvate (the substrates of the pathway) produce DXP.

Parameters from r10 (G3P + Pyruvate → DXP), catalyzed by DXS, are overrepresented in the top 51 highest sensitivities (14 out of 51), particularly *Vmax8* and *Km8* and for MEP pathway intermediates (Table 9). The highest absolute values of DXS-related sensitivities correspond to MEcPP sensitivities. This is consistent with experimental results showing that DXS activity is a determinant of MEcPP levels (Wright *et al*., 2014).

When we analyze the sensitivity of the positive fluxes to parameters, we find that 25 out of 728 sensitivities are above 0.5 in absolute value. Similarly, 21 out of 728 sensitivities of the negative fluxes are above 0.5 (Table 10). There is no specific parameter being overrepresented, and the pattern is that fluxes are most sensitive to Vmax and Km parameters from one of the cognate parameters of each flux. Interestingly, the highest sensitivities of the IPP/DMAPP input fluxes are to parameters from the IDI enzyme activity, not MVD or HDR.

**Table 10.** Logarithmic sensitivities of the aggregate flux to reaction parameters. Only sensitivities with an absolute value larger than 0.5 are shown.

| Positive flux |  |  | Negative flux |  |  |
| --- | --- | --- | --- | --- | --- |
| Metabolite | Parameter | Sensitivity | Metabolite | Parameter | Sensitivity |
| HMGC<sub>CoA</sub> | K<sub>i1</sub> | 0.923 | HMGC<sub>CoA</sub> | V<sub>max2</sub> | 1 |
| HMGC<sub>CoA</sub> | K<sub>m1</sub> | -0.931 | MVA | K<sub>m3</sub> | -1.000 |
| HMGC<sub>CoA</sub> | V<sub>max1</sub> | 1 | MVA | V<sub>max3</sub> | 1 |
| MVA | V<sub>max2</sub> | 1 | MVP | K<sub>m4</sub> | -0.999 |
| MVP | K<sub>m3</sub> | -1.000 | MVP | V<sub>max4</sub> | 1 |
| MVP | V<sub>max3</sub> | 1 | MVPP | K<sub>m5</sub> | -0.997 |
| MVPP | K<sub>m4</sub> | -0.999 | MVPP | V<sub>max5</sub> | 1 |
| MVPP | V<sub>max4</sub> | 1 | IPP<sub>cyt</sub> | k<sub>1</sub> | 1.000 |
| IPP<sub>cyt</sub> | V<sub>max7</sub> | 0.978 | IPP<sub>pla</sub> | k<sub>1'</sub> | 0.990 |
| IPP<sub>pla</sub> | V<sub>max7<sub>pl</sub></sub> | 0.988 | DMAPP<sub>cyt</sub> | k<sub>1</sub> | 1.000 |
| DMAPP<sub>cyt</sub> | K<sub>m6</sub> | -0.513 | DMAPP<sub>pla</sub> | k<sub>1'</sub> | 0.970 |
| DMAPP<sub>cyt</sub> | V<sub>max6</sub> | 0.955 | DXP | K<sub>m9</sub> | -0.917 |
| DMAPP<sub>pla</sub> | V<sub>max6<sub>pl</sub></sub> | 0.988 | DXP | V<sub>max9</sub> | 1 |
| DXP | K<sub>m8</sub> | -0.920 | MEP | K<sub>m10</sub> | -0.997 |
| DXP | V<sub>max8</sub> | 1 | MEP | V<sub>max10</sub> | 1 |
| MEP | K<sub>m9</sub> | -0.917 | CDPME | K<sub>m11</sub> | -0.999 |
| MEP | V<sub>max9</sub> | 1 | CDPME | V<sub>max11</sub> | 1 |
| CDPME | K<sub>m10</sub> | -0.997 | CDPMEP | K<sub>m12</sub> | -0.840 |
| CDPME | V<sub>max10</sub> | 1 | CDPMEP | V<sub>max12</sub> | 1 |
| CDPMEP | K<sub>m11</sub> | -0.999 | MECPP | K<sub>m13</sub> | -0.516 |
| CDPMEP | V<sub>max11</sub> | 1 | MECPP | V<sub>max13</sub> | 1 |
| MECPP | K<sub>m12</sub> | -0.840 |  |  |  |
| MECPP | V<sub>max12</sub> | 1 |  |  |  |
| HMBPP | K<sub>m13</sub> | -0.516 |  |  |  |
| HMBPP | V<sub>max13</sub> | 1 |  |  |  |

### 3.2 IPP and DMAPP levels are robust to circadian-dependent flux decrease

Model E implements regulatory effects of light on the biosynthesis of terpenoid precursors IPP and DMAPP. Thus, we analyze this model to understand how changes in the number of daylight hours affect IPP/DMAPP production.

First, we simulate the behavior of the system during two circadian oscillations. Figure 2 illustrates how the concentration of pathway intermediates and products increases through the day and decreases during the night, over 48 hours at different locations on the globe and in different seasons.

**Figure 2.**
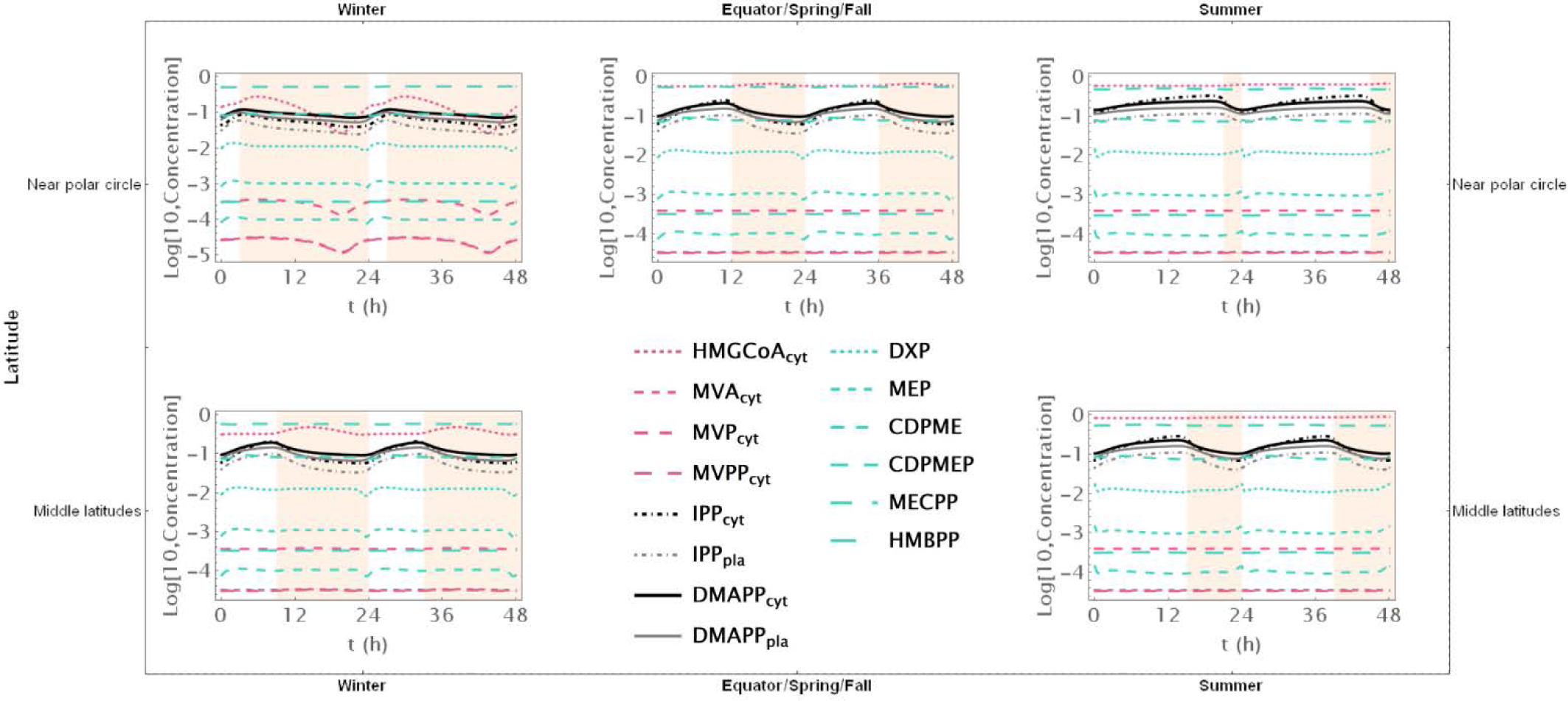
Time course simulation of the system throughout 48 h at different latitudes and times of the year: equator/spring and fall equinoxes (*dusk =* 12 h), middle latitudes (winter, *dusk =* 9 h; summer, *dusk* = 15 h) and near polar circle latitudes (winter, *dusk =* 3 h; summer, *dusk* = 21 h). *T* = 1 h.

When there are approximately 12 hours of daylight, the concentrations of intermediate metabolites have small oscillations in both pathways throughout the 24 hours of the day. This occurs consistently at or near the equator and in temperate latitudes during spring and fall. As the number of daylight hours per day increases, the relative amplitude of oscillation decreases. Similarly, as the number of daylight hours per day decreases, that amplitude increases. These changes are more pronounced for oscillations in the concentration of intermediate metabolites of the MVA pathway than for intermediates of the MEP pathway. The average daily concentrations of pathway intermediates is only slightly affected by the number of daylight hours. The picture is subtly reversed for the end products of the MVA and MEP pathways. The number of daylight hours per day has almost no influence in the relative amplitude of IPP and DMAPP concentration oscillations. In contrast, the daily average concentrations of IPP and DMAPP are directly correlated with the number of daylight hours.

Furthermore, the circadian rhythm significantly influences the system’s transient behavior. Cytosolic IPP and DMAPP steadily accumulate during daylight hours and decrease at night. In contrast, plastid IPP and DMAPP quickly reach quasi-steady state levels during daylight hours and gradually decrease at night. This behavior slightly changes during longer days, where quasi-steady states are reached for all four metabolite pools.

We further investigate how changing number of daylight hours between 0 and 24 affect IPP and DMAPP biosynthesis by simulating the dynamic behavior of the system when the number of daylight hours changes between 0 and 24. Then we measure the impact of those changes on concentration oscillations (Figures 3A-B) and production fluxes (Figures 3C, 3D, and 3E) within the pathways.

**Figure 3.**
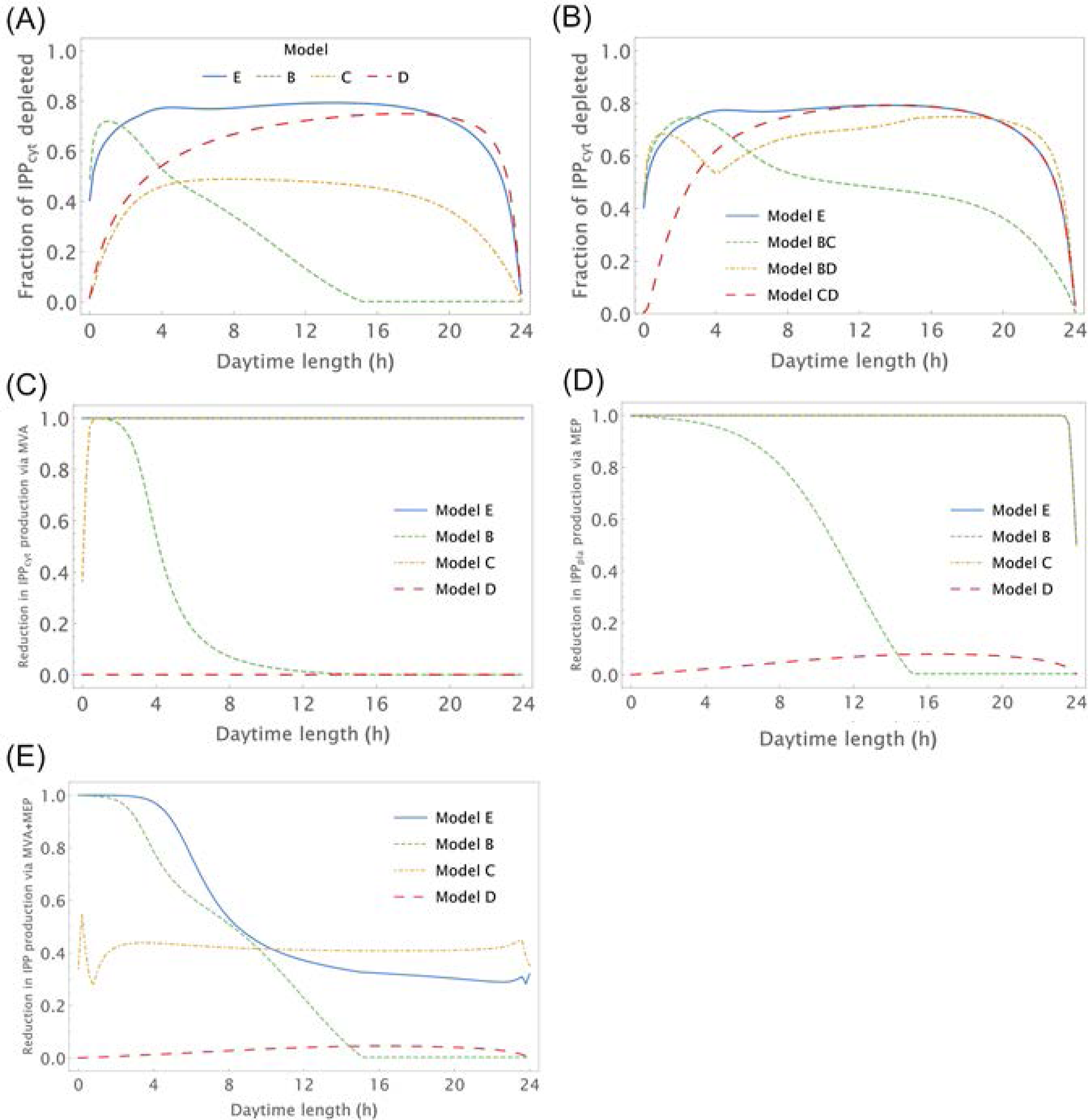
Relative amplitude of the circadian IPP concentration and flux oscillations as a function of the number of daylight hours (h). X-axis – number of daylight hours. Y-axis – minimum value/maximum value of a variable during a day. A, B) Relative amplitude of IPP_cyt_ concentration, C) Relative amplitude of the flux going through the MVA pathway to produce IPP, D) Relative amplitude of the flux going through the MEP pathway to produce IPP and E) Relative amplitude of the flux going through both pathwyas to produce IPP.

This experiment confirms that IPP concentration oscillations exhibit higher amplitudes than DMAPP (Figure 2). Additionally, we observe that the relative amplitude of IPP and DMAPP concentration oscillations remains relatively stable, except when the number of daylight hours approaches 0 or 24 (indicated by blue lines in Figure 3 panels), where the oscillation becomes a steady state with zero amplitude.

Overall, when plants experience between 3 and 21 daylight hours, the relative amplitude of IPP and DMAPP daily concentration oscillations remains roughly consistent (Figure 3A). The fluctuation in pathway-specific IPP production remains close to 100% regardless of daytime length (Figure 3C and 3D). However, overall production fluctuations can be significantly less than 100% if the daytime length exceeds 4 hours (Figure 3E), attributed to the anti-phasic regulation of pathway expression.

Essentially, while flux through one pathway nearly ceases at night, flux through the other pathway almost stops during daylight hours. When considering metabolite replenishment through isomerization and compartment exchange, production fluctuation is at most around 25%, even under short days (Figure S2).

### 3.3 The role of IPP-DMAPP exchange between the cytosol and the plastid

As mentioned in the introduction, while IPP and DMAPP are exchanged between the plastid and cytosol, there is little evidence that any other intermediate of either pathway also diffuses between compartments. Given that it is widely accepted that cytosolic and plastid IPP/DMAPP are used by the plant to synthesize quasi-orthogonal sets of more complex terpenoids, we were interested in understanding the effect of preventing the inter-compartmental diffusion of those metabolites. As such we removed that exchange from Model E, by setting diffusion rate parameters equal to zero.

This significantly decreases the amplitude of the IPP/DMAPP concentration oscillations induced by the circadian rhythm, while maintaining the amplitude of the oscillations at the level of pathway intermediates (Figure S3).

### 3.4 Contribution of the various regulatory modules towards the dynamic behavior of IPP and DMAPP biosynthesis during circadian light cycles

As described in section 2.7, three distinct regulatory modules connect the circadian rhythm to the regulation of MEP and MVA pathway activity. To assess the impact of these alternative regulatory modules, we investigate how the number of daylight hours per day influences the dynamics of IPP and DMAPP production in models B, C, D, BC, BD, and CD. We then replicate the analysis conducted for Model E.

#### 3.4.1 Circadian regulation of pathway substrate availability

Model B focuses solely on the circadian regulation of pathway substrate concentrations. Under this framework, IPP and DMAPP levels remain relatively stable, showing resilience to changes in daylight hours (refer to Figures 3A, and 4). Specifically, when exposed to more than 15 hours of daylight, IPP and DMAPP concentrations reach a quasi-steady state (as indicated by the green line in Figure 3). Oscillations become noticeable when daylight hours drop below 15, with the maximum relative amplitude of their concentration oscillations occurring at around 2 hours of daylight per day. Even under minimal daylight hours, slight oscillations still occur (as shown in Figure 4). However, the relative amplitude of concentration oscillations for pathway intermediates increases significantly compared to the fully regulated Model E (compare Figures 2 and 4). A noteworthy difference between Model E and Model B lies in the relative amplitude of concentrations for intermediates of the MVA and MEP pathways. Contrary to Model E, in Model B, the relative amplitude of concentrations for MVA pathway intermediates is smaller than that for MEP pathway intermediates.

**Figure 4.**
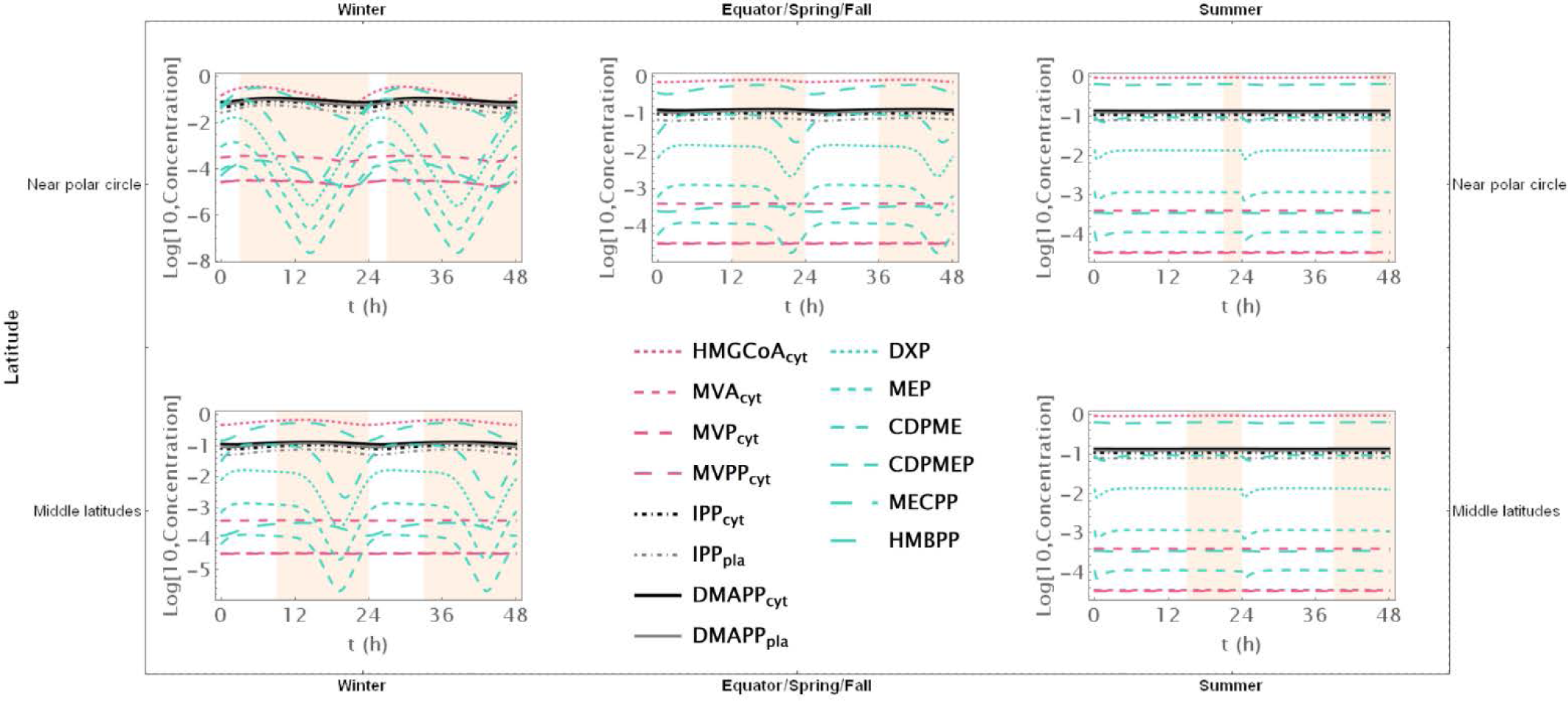
Model B. Time course simulation of the system throughout 48 h at different latitudes and times of the year: equator/spring and fall equinoxes (*dusk =* 12 h), middle latitudes (winter, *dusk =* 9 h; summer, *dusk* = 15 h) and near polar circle latitudes (winter, *dusk =* 3 h; summer, *dusk* = 21 h). *T* = 1 h.

Figure 3 illustrates the impact of varying daylight hours on the relative amplitude of IPP and DMAPP concentration oscillations. Days with more than 14 hours of daylight exhibit constant IPP and DMAPP concentrations. As daylight hours decrease, the effect of the circadian light rhythm on concentration oscillation amplitudes becomes similar to that observed in Model E (compare blue and green curves in Figure 3A). This shift results from significant decreases of up to 70% in metabolite concentrations during long nights compared to maximum daylight concentrations. It is also worth noting that Model B achieves nighttime IPP and DMAPP quasi-steady state concentrations in less than one hour while steadily accumulating these intermediates over daylight hours.

Circadian fluctuations in production via MVA and MEP pathways are observed on days shorter than 12 and 16 hours, respectively (refer to Figure 3C and 3D). Overall production of both pathways exhibits similar behavior (Figure 3E), and isomerization and compartment exchange dampen these oscillations.

#### 3.4.2 Antithetic circadian regulation of MEP and MVA gene expression

Model C only incorporates circadian regulation of gene expression in the MEP and MVA pathways, with an antithetic regulation pattern between the two pathways, as documented in previous studies (Atamian and Harmer, 2016; Covington *et al*., 2008; Vranová *et al*., 2013). Broadly, the behavior of IPP and DMAPP oscillations closely parallels that of model E: when plants experience between 3 and 21 daylight hours, the relative amplitude of daily concentration oscillations for IPP and DMAPP remains relatively stable. However, this amplitude is approximately half of that observed in Model E (compare blue and orange curves in Figure 3A). Additionally, a comparison between Figures 2 and 5 reveals that the dynamic behavior of pathway intermediate concentrations is similar between Models E and C. Moreover, variations in the fraction of daylight hours per day minimally affect these concentrations. Furthermore, variations in the fraction of daylight hours per day have minimal impact on these concentrations. In contrast, the relative amplitude of oscillations in the concentrations of pathway products IPP and DMAPP remains high when there are between 4 and 20 h of daylight.

**Figure 5.**
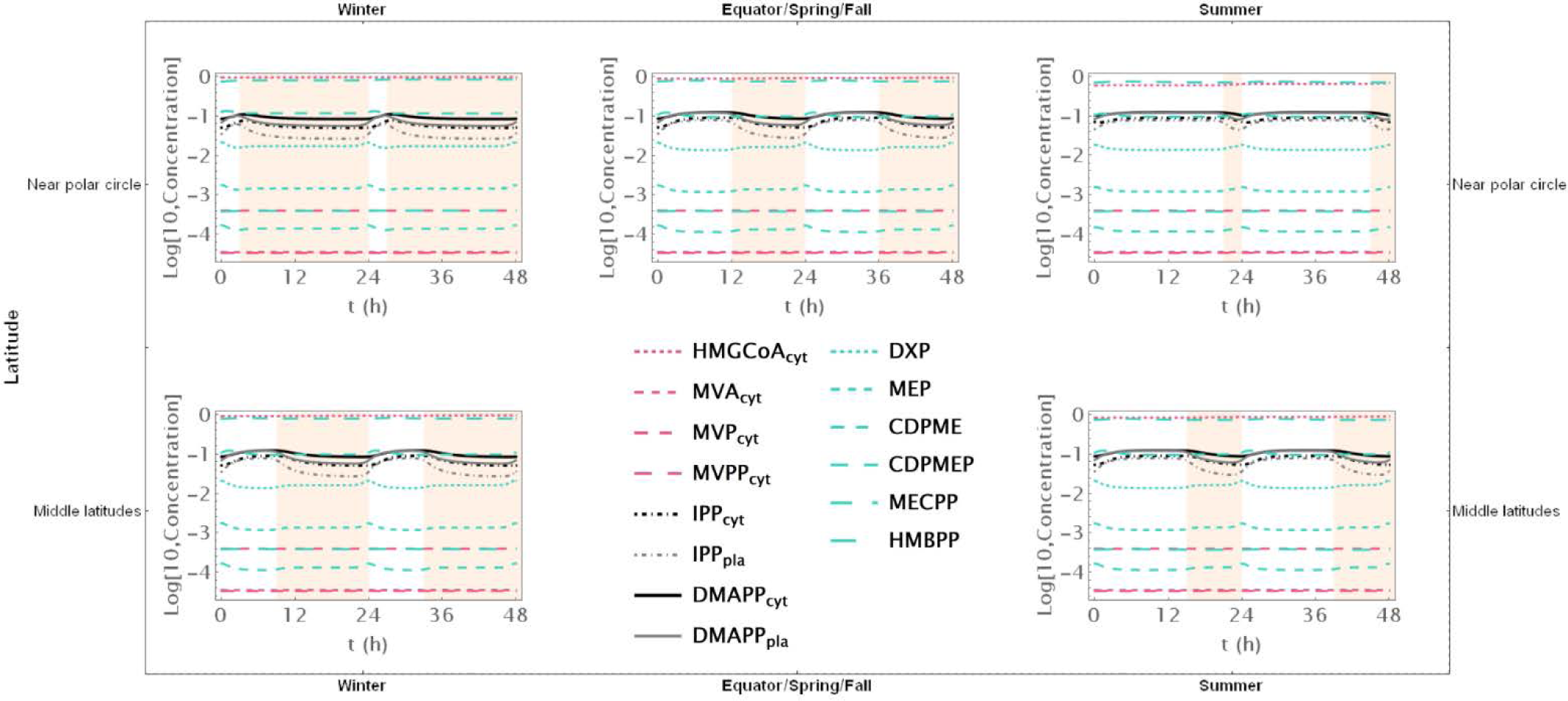
Model C. Time course simulation of the system throughout 48 h at different latitudes and times of the year: equator/spring and fall equinoxes (*dusk =* 12 h), middle latitudes (winter, *dusk =* 9 h; summer, *dusk* = 15 h) and near polar circle latitudes (winter, *dusk =* 3 h; summer, *dusk* = 21 h). *T* = 1 h.

Figure 3 demonstrates that the influence of daylight hours on the relative amplitude of the oscillations qualitatively mirrors that observed in model E. However, the depletion of IPP and DMAPP is approximately 20% smaller in this model compared to model E (compare orange and blue curves in Figure 3A). Interestingly, the system also demonstrates rapid adaptation to transitions between light and darkness, requiring approximately two hours to reach either a daylight quasi-steady state or a nighttime quasi-steady state. Model C also shows the same qualitative behavior as Model E in terms of flux circadian regulation. Global influx of IPP and DMAPP displays oscillations that are similar to those for concentration. The flux of material going through either the MVA or MEP pathways is reduced almost 100%, regardless of daytime length (Figure 3C and 3D). The aggregated fluxes show that the pathways take turns in producing IPP, as seen in Model E for long days (Figure 3E). In Model C, and because pathway substrate availability does not depend on the circadian rhythm, this type of dynamic behavior extends to short days as well.

#### 3.4.4 Circadian regulation of IPP and DMAPP consumption

Model D considers a situation where the circadian rhythm only regulates the activity of pathways that use IPP and DMAPP to synthesize more complex terpenoids. The red curve in Figure 3A shows that, under these conditions, the amplitude of IPP and DMAPP concentration oscillations undergoes sharp changes when the number of daylight hours is lower than 4. If the number of daylight hours is between 4 and 23, the oscillation amplitude experiences a slight increase. As the number of daylight hours goes up to 24, the oscillation amplitude diminishes, and the system reaches a steady state.

Similar to Model C, Figure 6 illustrates that pathway intermediate concentrations remain relatively stable across a broad range of daylight hours. Figure 6 shows that IPP and DMAPP concentrations oscillate in a pattern akin to that of Model E, while pathway intermediate levels maintain stability with low-amplitude oscillations, especially for MEP pathway intermediates. Throughout the day, both compartments witness accumulation of IPP and DMAPP, followed by rapid depletion at night. In Model D, maximum relative depletions occur for days with 23 hours of light.

**Figure 6.**
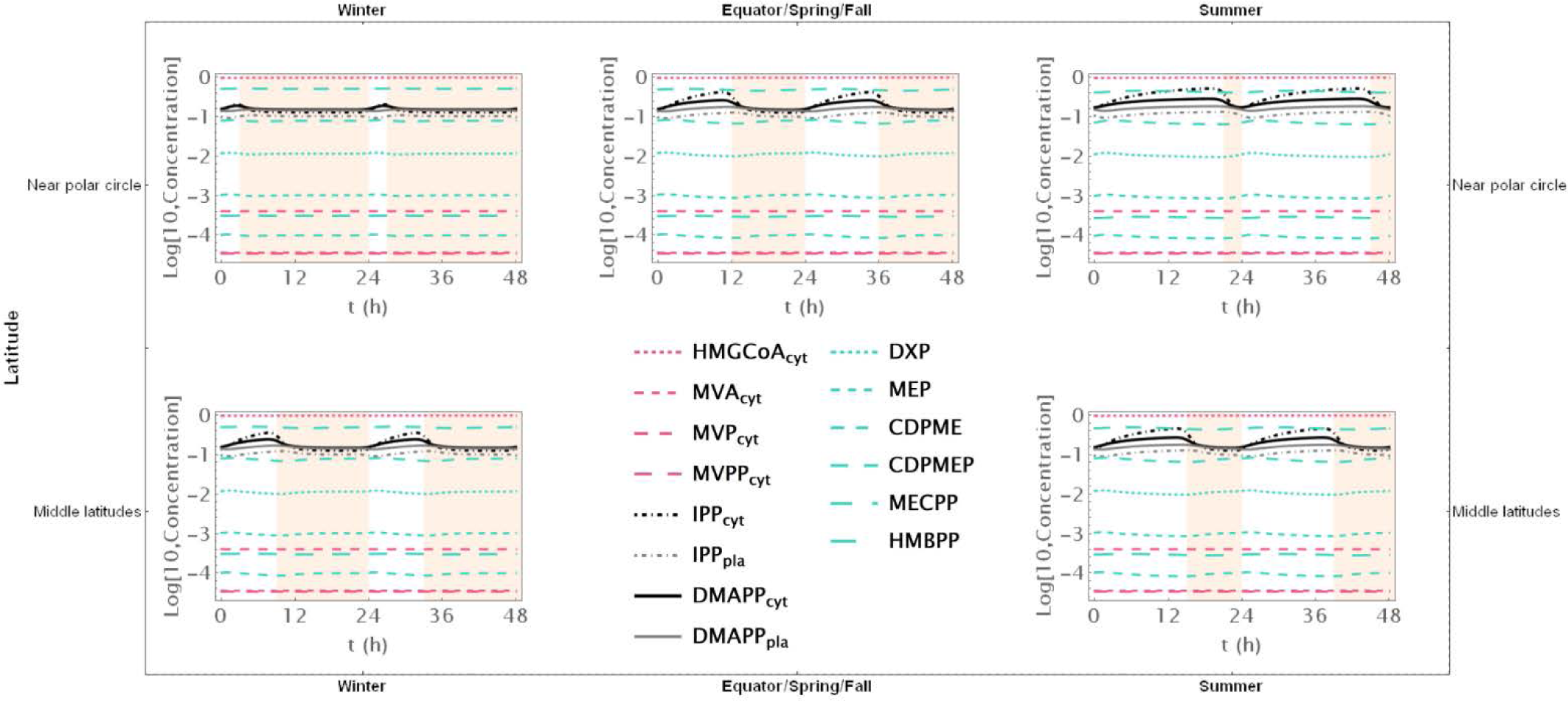
Model D. Time course simulation of the system throughout 48 h at different latitudes and times of the year: equator/spring and fall equinoxes (*dusk =* 12 h), middle latitudes (winter, *dusk =* 9 h; summer, *dusk* = 15 h) and near polar circle latitudes (winter, *dusk =* 3 h; summer, *dusk* = 21 h). *T* = 1 h.

The flux of material going through the MVA pathway is approximately constant regardless of daytime length (Figure 3C). The flux of material going through the MEP pathway is reduced, at most, by 10% (see the behavior in Figure 3D). The same behavior is observed for the sum of both fluxes, but with slightly weaker oscillations (Figure 3E). Maximum oscillation amplitude happens at daytime length around 18h.

#### 3.3.5 Pairwise combination of circadian regulatory modules

Models BC, CD, and BD simulate situations where part of the circadian regulation is lost. In Model BC circadian regulation of substrate availability is lost. In Model BD circadian regulation of gene expression is lost. In Model CD circadian regulation of IPP/DMAPP utilization is lost. We apply the same analysis as with the previous models (Figure 3E). In general, the relationship between the amplitude of concentration oscillations and the number of daylight hours in a model with two active circadian regulation modules resembles the combined dependencies of models where each individual module is the sole active circadian regulator (compare Figures 3A and 3B). In addition, the amplitude of concentration and flux oscillations is always smaller than that observed for the fully regulated Model E.

## 4 Discussion

### 4.1 Cellular demand for IPP/DMAPP and the MVA and MEP pathways

IPP and DMAPP are the precursor monomers to terpenoids, a family of molecules that contains many chemicals with importance in biology, pharmacy, biotechnology, biomedicine and cosmetics, such as squalene, cholesterol, some vitamins and most plant hormones. Plants produce those monomers using two biosynthetic pathways: the MVA pathway in the cytosol, and the MEP pathway in the plastid.

IPP and DMAPP are used as the building blocks for more complex terpenoids, ranging from protective molecules such as carotenoids to hormones such as strigolactones. A version of the MVA pathway was present in the last common ancestor of archaea and eukaryotes, while ancestral bacteria contained a version of the MEP pathway (Hoshino and Gaucher, 2018; Zeng and Dehesh, 2021; Lombard and Moreira, 2011). In plants, the current MEP pathway seems to have evolved from the ancestral MEP pathway present in the early symbiotic cyanobacteria that became the chloroplast (Lichtenthaler, 1999). While both pathways have a level of crosstalk and IPP and DMAPP can be exchanged between the cytosol and the plastid, the contribution of each pathway to the biosynthesis of complex terpenoids is not the same. The MVA pathway mainly provides flux for the biosynthesis of sesquiterpenes, sterols, polyprenols, and triterpenes, while the MEP pathway is the main provider for the biosynthesis of chlorophylls, tocopherols, quinones, carotenoids, monoterpenes and strigolactones, among others (Pérez *et al*., 2022). Both pathways provide flux and materials for cellular metabolism. As such one would expect that their regulation would be mainly demand-driven, that is, its flux should be mainly regulated by the cellular demand for the material produced by the pathways.

### 4.2 Regulatory design for the MVA and MEP pathways is consistent with design principles for demand driven pathways

Several hallmarks of demand-driven pathways exist (Savageau, 1972; R. Alves and Savageau, 2000; Bromig *et al*., 2020). In terms of regulation, demand-driven pathways are more efficiently regulated by negative feedback (R. Alves and Savageau, 2000; Savageau, 1972). The most efficient negative feedback configuration is created by overall feedback, where the product of the pathway inhibits the flux of the first reaction (Alves and Savageau, 2001; R. Alves and Savageau, 2000; Ye and Medzhitov, 2019; Savageau, 1972). This is clearly the case with the MEP pathway (Figure 1).

Intriguingly, this type of regulation is absent from the MVA pathway. One can speculate why this is so. A probable reason for the absence of overall feedback in the MVA pathway is that the synthesis of the final products of the cytosolic MVA pathway, IPP and DMAPP, occurs in the peroxisome, not in the cytosol (Vranová *et al*., 2013). This prevents overall negative feedback from these products to the initial step of the pathway. As such, the cascading inhibitory feedback observed for the MVA pathway, where each enzyme is inhibited by its own product, still conveys information about cellular demand for the end product backwards through the biosynthetic chain one step at a time. This feedback structure creates a pathway whose response time to changes in cellular demand is slower than that permitted by the overall feedback configuration (R. Alves and Savageau, 2000), but still responsive to cellular demand. Another signature behavior for demand-driven pathways is that the concentration of pathway intermediates is lower than that of the pathway final products. This is observed when the steady state of Model E is calculated (Table 6). Stability (Table 7) and robustness of the steady state to changes in the parameter values (Tables 9 and 10) are other signature of demand-driven pathways (R. Alves and Savageau, 2000; Savageau, 1972). Both pathways have stable steady states when subjected to constant light conditions, and that steady state is robust to change in model parameters. Additionally, we find that IPP and DMAPP levels are less sensitive to restrictions on carbohydrate availability than the levels of pathway intermediates.

### 4.3 Circadian regulation of supply, gene expression and demand for the MVA and MEP pathways

The circadian light cycle regulates MVA and MEP pathway activity at three levels: pathway substrate availability, expression of genes coding for pathway enzymes, and activity of pathways that use IPP and DMAPP for the synthesis of more complex terpenoids. Figure 2 shows that some of the steady state design principles described in section 4.2 are also observed when pathway dynamic behavior is away from the steady state and driven by the circadian light cycle. On the one hand, IPP and DMAPP concentrations are orders of magnitude higher than those for most pathway intermediates for both the MVA and the MEP pathways (Figure 2). On the other hand, the metabolic oscillation itself is insensitive over a wide range to changes between shorter and longer days. For days with between 4 and 20 hours of light, the amplitude of the oscillation remains surprisingly stable (Figure 3).

Understanding how each of the three regulatory levels contributes to that stability drove us to create models of the pathways were circadian regulation was only active for one of the levels. When the light cycle only regulates pathway substrate availability, we find that the concentration waves for pathway intermediates are much more sensitive to changes in the number of daylight hours in the circadian rhythm than for the fully regulated model (Figure 4). However, the concentration of the products IPP and DMAPP remains very insensitive to the supply of pathway substrate. In fact, if the number of daylight hours is above 15, the oscillation is lost, and the pathway operates at or near a steady state. This further strengthens the argument that the structure and parameters of these pathways have been selected by evolution to be consistent with a demand driven pathway. Our model shows that MVA-produced IPP and DMAPP oscillate with smaller relative amplitudes than those for the concentrations of the intermediate metabolites of the pathway. The relative amplitude of the IPP and DMAPP concentration oscillations is approximately half of that observed when the circadian rhythm is considered to regulate the three levels of pathway activity, while remaining equally insensitive to changes in the number of daylight hours.

When the light cycle only regulates gene expression, concentrations of pathway intermediates are very insensitive to changes in the number of daylight hours (Figure 5). Posttranscriptional regulation is important for the proper functioning of the MEP and MVA pathways in plants (Guevara-García *et al*., 2005; Flores-Pérez *et al*., 2008; Sauret-Güeto *et al*., 2006; Cordoba *et al*., 2009; Xie *et al*., 2008; Han *et al*., 2013; Laule *et al*., 2003). When this is the only type of circadian regulation acting on the MVA and MEP pathways, our models indicate that IPP/DMAPP concentrations are significantly less robust to fluctuations in enzyme activity than the concentrations of pathway intermediates. As such, circadian regulation of gene expression alone, would create a regulatory structure for the pathway that would be suboptimal for regulating a demand driven pathway. We also found that having antithetic regulation of the gene expression between the MVA and the MEP pathways leads to oscillations that have smaller amplitudes, making the dynamic flux going through the pathways less sensitive to changes between day and night.

When the light cycle only regulates IPP and DMAPP utilization, concentrations of pathway intermediates remain insensitive to changes in the number of daylight hours (Figure 6). The amplitude of the IPP and DMAPP concentration oscillations is similar to that observed when the circadian rhythm is considered to regulate the three levels of pathway activity. However, it is more sensitive to changes between longer and shorter days, making the concentrations and fluxes for the system very responsive to changes in cellular demand for IPP and DMAPP.

Taken together, these results suggest that the three levels of circadian regulation, plus the MVA- and MEP-specific regulatory inhibition loops contribute differently to creating an operating regime that maintains pathway flux strongly coupled to demand and insensitive to changes over a wide range of daylight hours. Inhibitory feedback stabilizes the pathway product concentrations, when circadian rhythms change pathway supply availability, at the cost of amplifying concentration oscillations of pathway intermediates. While circadian regulation acts only on gene expression or on demand for IPP/DMAPP, the same inhibitory feedback creates product concentration oscillations with bigger amplitudes and decreases the amplitude of the oscillations for pathway intermediates. Thus, circadian regulation of gene expression or of demand for IPP/DMAPP alone would create a pathway whose dynamic response is suboptimal to demand for the pathways’ final products. It is the joining of the three regulatory modules that balances the dynamic behavior of the pathway, making it as robust as possible to the cellular demand for pathway products.

### 4.4 Limitations of this work

Here we present what we believe are the main limitations of this study and discuss how they could affect the robustness and applicability of its findings.

One important limitation of this study is the black box manner in which we model the production of substrate for the MVA and MEP pathways. This is a limitation that is shared with other modeling studies (Rios-Estepa *et al*., 2010; Pokhilko *et al*., 2015; Neiburga *et al*., 2023; Weaver *et al*., 2015; Basallo *et al*., 2023), and we believe that this valid simplification facilitates the analysis of the dynamic and regulatory behavior of the MVA and MEP pathways.

Another limitation is the fact that we use a similar approach to model the consumption of IPP and DMAPP out of the two pathways for the biosynthesis of more complex terpenoids. The biosynthesis of these and other terpenoid final products is also treated in this paper as a black box that draws flux from the MEP and MVA pathways in a way that is dependent on the circadian rhythm. This is valid for many volatile terpenoids, as well as carotenoids and other phytohormones (Loivamäki *et al*., 2007; Zheng *et al*., 2017; Zeng *et al*., 2017; Mu *et al*., 2022; Picazo-Aragonés *et al*., 2020; Covington *et al*., 2008). Still, to more accurately understand how the biosynthesis of these more complex terpenoids affects the dynamics of the MEP and MVA pathways, additional research is needed. This research should develop, analyze, and integrate detailed models of the biosynthetic pathways for those terpenoids with the MVA and MEP pathway models. That development and integration should take into account that certain terpenoids are synthesized using material that is drawn mainly from only one of the pathways (Qiao *et al*., 2021; Yang *et al*., 2012; Rodríguez-Concepción *et al*., 2004; Wu *et al*., 2021; Chandrasekaran *et al*., 2022; Liu *et al*., 2024). MVA-derived isoprenoid end products in plants are sterols and cytokinins that modulate membrane architecture, plant growth and development, and brassinosteroids that work as steroid hormones. In contrast, MEP-derived end products include photosynthesis-related isoprenoids (carotenoids and the side chains of chlorophylls, plastoquinones, and phylloquinones), gibberellins and abscisic acid hormones, and root volatile monoterpenes.

An additional limitation of this work is that the sensitivity analysis we performed, while comprehensive in identifying parameters to which the model is highly sensitive, is differential. This approach may not fully capture the dynamic complexities of biological systems and may miss higher order interactions between simultaneous changes in more than one parameter. We also note that, while the study provides insights into how different regulatory modules influence the effect of the circadian rhythm and latitude on IPP and DMAPP production, it does not fully address the potential interactions between these modules and other environmental or physiological factors that might affect the circadian rhythm. The models used to simulate these regulatory effects (Models B, C, D, and their combinations) are simplified representations and may not capture all the nuances of circadian regulation in real biological systems. For example, the antithetic regulation of gene expression in Model C and the regulation of consumption pathways in Model D may not fully represent the intricate feedback mechanisms present *in vivo*.

### 4.5 Conclusions

Several conclusions can be drawn from this work. Because the dynamic behavior of our model is robust, this allows us to conclude that its use to simulate physiological situations is likely to be appropriate. Our analysis also concludes that the feedback inhibition of enzymes by pathway intermediates and end products is compatible with a situation where the regulation of the flux going through those pathways is significantly driven by the cellular demand for their end product. Finally, we can also conclude that the three regulatory modules at which the circadian rhythm affects IPP and DMAPP production interact to make that production less sensitive to the seasonal changes in the number of daylight hours observed at different latitudes in our planet. Finally, the antithetic regulation in gene expression contributes to buffer the global production of IPP and DMAPP against shifts between day and night.

Thus, even with very limited quantitative information available, mathematical models can point at relevant features of these pathway and propose scenarios for future experimental exploration that may facilitate the modification of IPP and DMAPP production through synthetic biology efforts.

## 8 Data Availability Statement

The datasets and scripts generated for this study can be found in Supplementary Dataset1.

---

**Figure S1.**
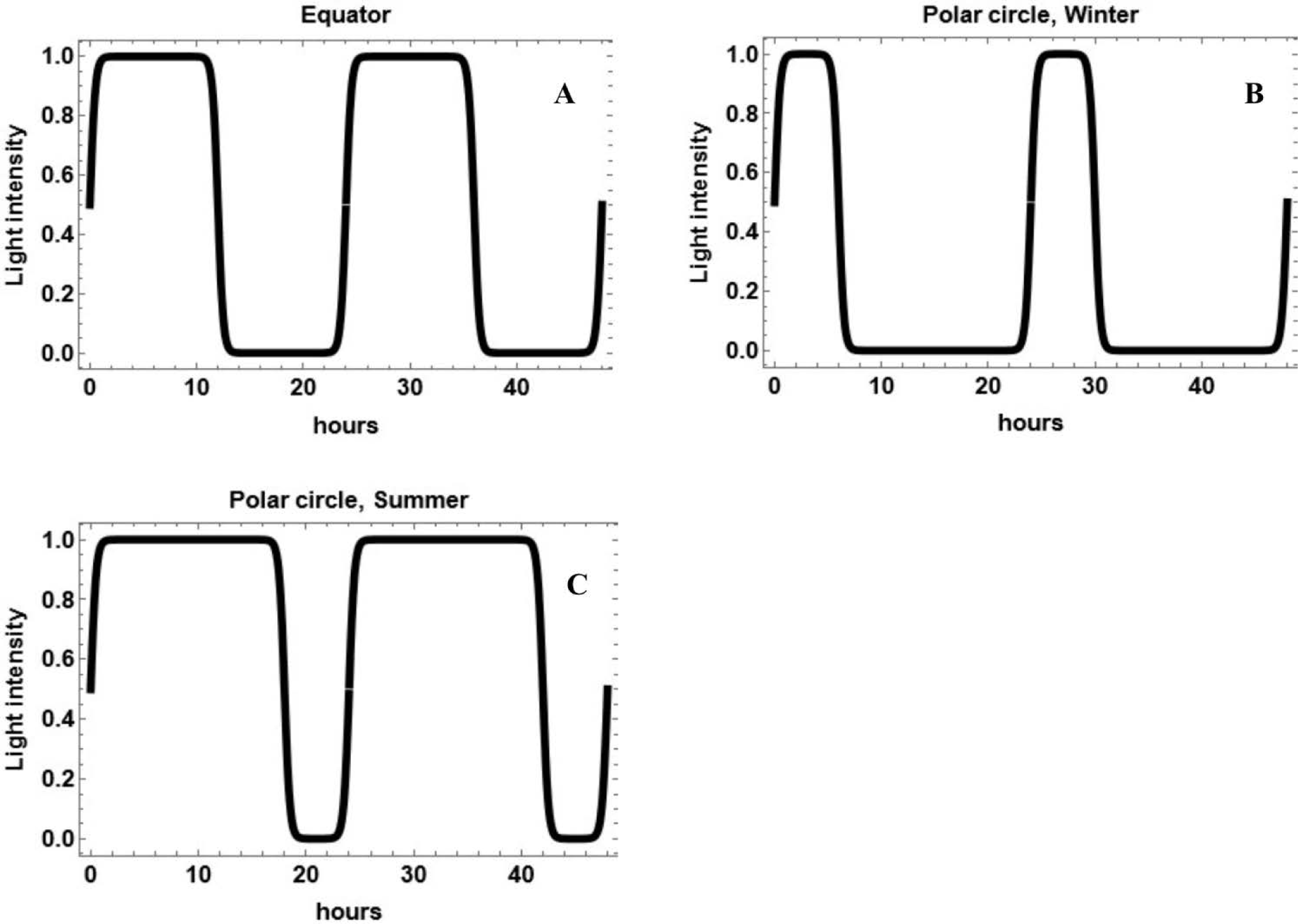
Modeling circadian light cycles with the L function. A – Approximate daylight hours at the equator. B – Approximate daylight hours at the polar circle during the peak of winter. C – Approximate daylight hours at the polar circle during the peak of summer.

**Figure S2.**
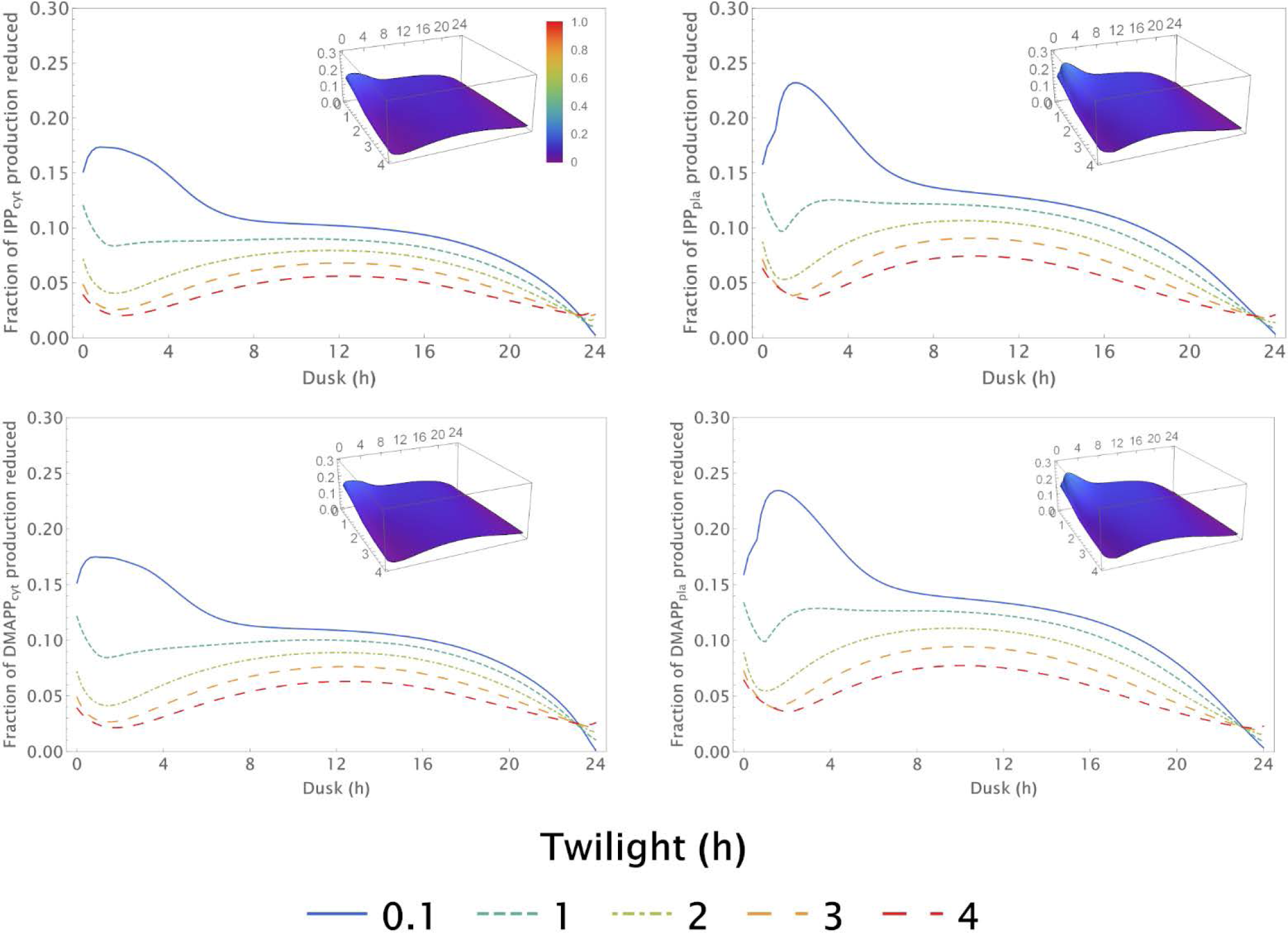
Model E. Oscillation amplitude of IPP and DMAPP production (normalized to the maxima) for different values of *T* (Twilight) and different values of *dusk* (Daytime).

**Figure S3.**
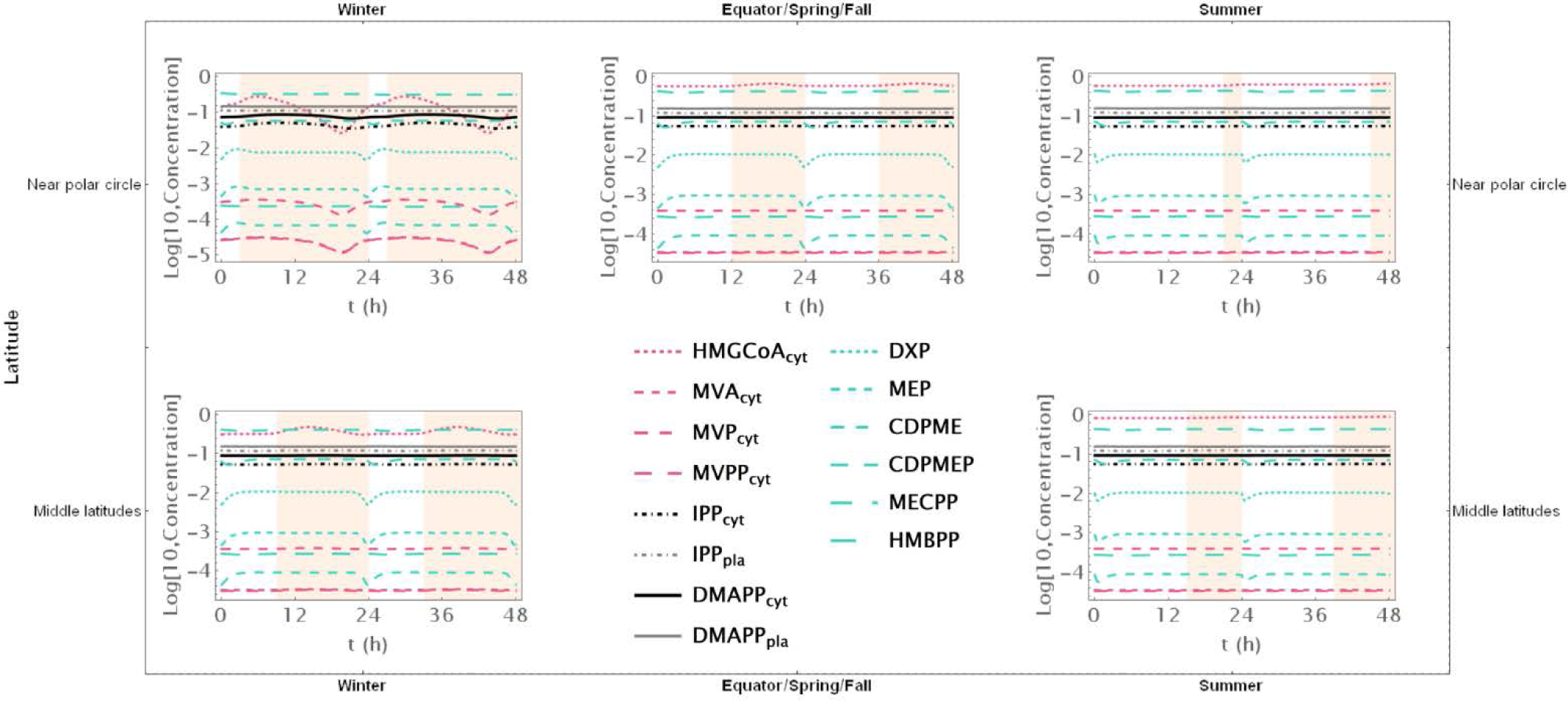
Model E without diffusion of IPP/DMAPP between compartments. Time course simulation of the system throughout 48 h at different latitudes and times of the year: equator/spring and fall equinoxes (*dusk =* 12 h), middle latitudes (winter, *dusk =* 9 h; summer, *dusk* = 15 h) and near polar circle latitudes (winter, *dusk =* 3 h; summer, *dusk* = 21 h). *T* = 1 h.

## References

Alabadí, D. et al. (2001) Reciprocal regulation between TOC1 and LHY/CCA1 within the Arabidopsis circadian clock. Science, 293, 880–883.

Albe, K.R. et al. (1990) Cellular Concentrations of Enzymes and Their Substrates.

Alves, R. et al. (2008) Mathematical formalisms based on approximated kinetic representations for modeling genetic and metabolic pathways. Biotechnol. Genet. Eng. Rev., 25, 1–40.

Alves, Rui and Savageau, M.A. (2000) Comparing systemic properties of ensembles of biological networks by graphical and statistical methods. Bioinformatics, 16, 527–533.

Alves, R. and Savageau, M.A. (2000) Effect of overall feedback inhibition in unbranched biosynthetic pathways. Biophys. J., 79, 2290–2304.

Alves, R. and Savageau, M.A. (2001) Irreversibility in unbranched pathways: preferred positions based on regulatory considerations. Biophys. J., 80, 1174–1185.

Atamian, H.S. and Harmer, S.L. (2016) Circadian regulation of hormone signaling and plant physiology. Plant Mol. Biol. 2016 916, 91, 691–702.

Baadhe, R.R. et al. (2012) Development of petri net-based dynamic model for improved production of farnesyl pyrophosphate by integrating mevalonate and methylerythritol phosphate pathways in yeast. Appl. Biochem. Biotechnol., 167, 1172–1182.

Basallo, O. et al. (2023) Changing biosynthesis of terpenoid percursors in rice through synthetic biology. Front. Plant Sci., 14, 1133299.

Baskaran, G. et al. (2015) HMG-CoA reductase inhibitory activity and phytocomponent investigation of Basella alba leaf extract as a treatment for hypercholesterolemia. Drug Des. Devel. Ther., 9, 509–517.

Bick, J.A. and Lange, B.M. (2003) Metabolic cross talk between cytosolic and plastidial pathways of isoprenoid biosynthesis: unidirectional transport of intermediates across the chloroplast envelope membrane. Arch. Biochem. Biophys., 415, 146–154.

BRENDA Enzyme Database (2024).

Bromig, L. et al. (2020) Understanding biochemical design principles with ensembles of canonical non-linear models. PloS One, 15, e0230599.

Buchanan, B.B., et al. (2002) Biochemistry & molecular biology of plants Buchanan, B.B. et al. (eds) American Society of Plant Physiologists.

Chandrasekaran, U. et al. (2022) Short-term severe drought influences root volatile biosynthesis in eastern white pine (Pinus strobus L). Front. Plant Sci., 13.

Cockburn, W. and McAulay, A. (1977) Changes in Metabolite Levels in Kalanchoë daigremontiana and the Regulation of Malic Acid Accumulation in Crassulacean Acid Metabolism. Plant Physiol., 59, 455–458.

Cordoba, E. et al. (2009) Unravelling the regulatory mechanisms that modulate the MEP pathway in higher plants. J. Exp. Bot., 60, 2933–2943.

Covington, M.F. et al. (2008) Global transcriptome analysis reveals circadian regulation of key pathways in plant growth and development. Genome Biol., 9, 1–18.

Dudareva, N. et al. (2005) The nonmevalonate pathway supports both monoterpene and sesquiterpene formation in snapdragon flowers. Proc. Natl. Acad. Sci. U. S. A., 102, 933–938.

Eisenreich, W. et al. (2001) Deoxyxylulose phosphate pathway to terpenoids. Trends Plant Sci., 6, 78–84.

Flores-Pérez, Ú. et al. (2008) A Mutant Impaired in the Production of Plastome-Encoded Proteins Uncovers a Mechanism for the Homeostasis of Isoprenoid Biosynthetic Enzymes in Arabidopsis Plastids. Plant Cell, 20, 1303–1315.

Guevara-García, A. et al. (2005) Characterization of the Arabidopsis clb6 Mutant Illustrates the Importance of Posttranscriptional Regulation of the Methyl-d-Erythritol 4-Phosphate Pathway. Plant Cell, 17, 628–643.

Han, M. et al. (2013) Enzyme inhibitor studies reveal complex control of methyl-D-erythritol 4- phosphate (MEP) pathway enzyme expression in Catharanthus roseus. PloS One, 8.

Harborne, J.B. (Jeffrey B.) et al. (1991) Ecological chemistry and biochemistry of plant terpenoids Clarendon Press.

Hemmerlin, A. et al. (2006) A Cytosolic Arabidopsis d-Xylulose Kinase Catalyzes the Phosphorylation of 1-Deoxy-d-Xylulose into a Precursor of the Plastidial Isoprenoid Pathway. Plant Physiol., 142, 441–457.

Hemmerlin, A. et al. (2012) A raison d’être for two distinct pathways in the early steps of plant isoprenoid biosynthesis? Prog. Lipid Res., 51, 95–148.

Hemmerlin, A. et al. (2003) Cross-talk between the cytosolic mevalonate and the plastidial methylerythritol phosphate pathways in tobacco bright yellow-2 cells. J. Biol. Chem., 278, 26666–26676.

Hemmerlin, A. (2013) Post-translational events and modifications regulating plant enzymes involved in isoprenoid precursor biosynthesis. Plant Sci., **203–204**, 41–54.

Hoshino, Y. and Gaucher, E.A. (2018) On the Origin of Isoprenoid Biosynthesis. Mol. Biol. Evol., 35, 2185–2197.

Jin, X. et al. (2021) The Coordinated Upregulated Expression of Genes Involved in MEP, Chlorophyll, Carotenoid and Tocopherol Pathways, Mirrored the Corresponding Metabolite Contents in Rice Leaves during De-Etiolation. Plants, 10, 1456.

Kitano, H. (2007) Towards a theory of biological robustness. Mol. Syst. Biol., 3, 137.

Lange, I. et al. (2015) Comprehensive Assessment of Transcriptional Regulation Facilitates Metabolic Engineering of Isoprenoid Accumulation in Arabidopsis. Plant Physiol., 169, 1595–1606.

Laule, O. et al. (2003) Crosstalk between cytosolic and plastidial pathways of isoprenoid biosynthesis in Arabidopsis thaliana. Proc. Natl. Acad. Sci. U. S. A., 100, 6866–71.

Lee, C.P. et al. (2021) The versatility of plant organic acid metabolism in leaves is underpinned by mitochondrial malate–citrate exchange. Plant Cell, 33, 3700.

Liao, P. et al. (2016) The potential of the mevalonate pathway for enhanced isoprenoid production. Biotechnol. Adv., 34, 697–713.

Lichtenthaler, H.K. (1999) The 1-Deoxy-D-Xylulose-5-Phosphate Pathway Of Isoprenoid Biosynthesis In Plants. Annu. Rev. Plant Physiol. Plant Mol. Biol., 50, 47–65.

Liu, J. et al. (2024) Classification, biosynthesis, and biological functions of triterpene esters in plants. Plant Commun., 5, 100845.

Loivamäki, M. et al. (2007) Circadian Rhythms of Isoprene Biosynthesis in Grey Poplar Leaves. Plant Physiol., 143, 540–551.

Lombard, J. and Moreira, D. (2011) Origins and Early Evolution of the Mevalonate Pathway of Isoprenoid Biosynthesis in the Three Domains of Life. Mol. Biol. Evol., 28, 87–99.

McClung, C.R. (2013) Beyond Arabidopsis: The circadian clock in non-model plant species. Semin. Cell Dev. Biol., 24, 430–436.

Mcgarvey, D.J. and Croteau, R. (1995) Terpenoid Metabolism American Society of Plant Physiologists.

Mu, Z. et al. (2022) An Overview of the Isoprenoid Emissions From Tropical Plant Species. Front. Plant Sci., 13, 833030.

Nagel, D.H. and Kay, S.A. (2012) Complexity in the Wiring and Regulation of Plant Circadian Networks. Curr. Biol., 22, R648–R657.

Neiburga, K.D. et al. (2023) Total optimization potential (TOP) approach based constrained design of isoprene and cis-abienol production in A. thaliana. Biochem. Eng. J., 190, 108723.

Page, J.E. et al. (2004) Functional Analysis of the Final Steps of the 1-Deoxy-d-xylulose 5-phosphate (DXP) Pathway to Isoprenoids in Plants Using Virus-Induced Gene Silencing. Plant Physiol., 134, 1401–1413.

Pérez, L. et al. (2022) Multilevel interactions between native and ectopic isoprenoid pathways affect global metabolism in rice. Transgenic Res., 31, 249–268.

Picazo-Aragonés, J. et al. (2020) Plant Volatile Organic Compounds Evolution: Transcriptional Regulation, Epigenetics and Polyploidy. Int. J. Mol. Sci. 2020 Vol 21 Page 8956, **21**, 8956.

Pokhilko, A. et al. (2014) Adjustment of carbon fluxes to light conditions regulates the daily turnover of starch in plants: a computational model. Mol. Biosyst., 10, 613–627.

Pokhilko, A. et al. (2015) Mathematical modelling of the diurnal regulation of the MEP pathway in Arabidopsis. New Phytol., 206, 1075–1085.

Qiao, Z. et al. (2021) An Update on the Function, Biosynthesis and Regulation of Floral Volatile Terpenoids. Horticulturae, 7, 451.

Rios-Estepa, R. et al. (2010) Mathematical modeling-guided evaluation of biochemical, developmental, environmental, and genotypic determinants of essential oil composition and yield in peppermint leaves. Plant Physiol., 152, 2105–2119.

Rodríguez-Concepción, M. et al. (2004) Distinct Light-Mediated Pathways Regulate the Biosynthesis and Exchange of Isoprenoid Precursors during Arabidopsis Seedling Development. Plant Cell, 16, 144–156.

Sauret-Güeto, S. et al. (2006) Plastid cues posttranscriptionally regulate the accumulation of key enzymes of the methylerythritol phosphate pathway in Arabidopsis. Plant Physiol., 141, 75– 84.

Savageau, M.A. (1976) Biochemical systems analysis : a study of function and design in molecular biology. 379.

Savageau, M.A. (1972) The Behavior of Intact Biochemical Control Systems*. In, Horecker, B.L. and Stadtman, E.R. (eds), Current Topics in Cellular Regulation. Academic Press, pp. 63–130.

Schaller, H. et al. (1995) Expression of the Hevea brasiliensis (H.B.K.) Mull. Arg. 3-Hydroxy-3- Methylglutaryl-Coenzyme A Reductase 1 in Tobacco Results in Sterol Overproduction. Plant Physiol., **109**, 761–770.

Singh, V.K. and Ghosh, I. (2013) Methylerythritol phosphate pathway to isoprenoids: Kinetic modeling and in silico enzyme inhibitions in Plasmodium falciparum. FEBS Lett., 587, 2806– 2817.

Sorribas, A. et al. (2007) Cooperativity and saturation in biochemical networks: a saturable formalism using Taylor series approximations. Biotechnol. Bioeng., 97, 1259–1277.

Taiz, L., et al. (2014) Plant Physiology and Development 6th ed. 20. Taiz, L. et al. (eds) Sinauer.

Tetali, S.D. (2019) Terpenes and isoprenoids: a wealth of compounds for global use. Planta, 249, 1–8.

Voit, E.O. (2013) Biochemical Systems Theory: A Review. ISRN Biomath., 2013, 1–53.

Voit, E.O. (1991) Canonical nonlinear modeling : S-system approach to understanding complexity. 365.

Vranová, E., et al. (2013) Network Analysis of the MVA and MEP Pathways for Isoprenoid Synthesis. Httpsdoiorg 101146annurev-Arplant-050312-120116, 64, 665–700.

Weaver, L.J. et al. (2015) A kinetic-based approach to understanding heterologous mevalonate pathway function in E. coli. Biotechnol. Bioeng., 112, 111–119.

Wishart, D.S. et al. (2022) HMDB 5.0: the Human Metabolome Database for 2022. Nucleic Acids Res., 50, D622–D631.

Wright, L.P. et al. (2014) Deoxyxylulose 5-Phosphate Synthase Controls Flux through the Methylerythritol 4-Phosphate Pathway in Arabidopsis. Plant Physiol., 165, 1488–1504.

Wu, W. et al. (2021) The diverse roles of cytokinins in regulating leaf development. Hortic. Res., 8, 1–13.

Xie, Z. et al. (2008) A systems biology investigation of the MEP/terpenoid and shikimate/phenylpropanoid pathways points to multiple levels of metabolic control in sweet basil glandular trichomes. Plant J., 54, 349–361.

Yang, D. et al. (2012) Different Roles of the Mevalonate and Methylerythritol Phosphate Pathways in Cell Growth and Tanshinone Production of Salvia miltiorrhiza Hairy Roots. PLOS ONE, 7, e46797.

Ye, J. and Medzhitov, R. (2019) Control strategies in systemic metabolism. Nat. Metab., 1, 947–957.

Yon, F. et al. (2017) Fitness consequences of altering floral circadian oscillations for Nicotiana attenuata. J. Integr. Plant Biol., 59, 180–189.

Zeng, L. et al. (2017) Regulation of the Rhythmic Emission of Plant Volatiles by the Circadian Clock. Int. J. Mol. Sci. 2017 *Vol 18 Page* 2408, **18**, 2408.

Zeng, L. and Dehesh, K. (2021) The eukaryotic MEP-pathway genes are evolutionarily conserved and originated from Chlaymidia and cyanobacteria. BMC Genomics, 22, 137.

Zheng, R. et al. (2017) Expression of mep pathway genes and non-volatile sequestration are associated with circadian rhythm of dominant terpenoids emission in osmanthus fragrans lour. Flowers. Front. Plant Sci., 8, 259087.

Zhou, F. and Pichersky, E. (2020) More is better: the diversity of terpene metabolism in plants. Curr. Opin. Plant Biol., 55, 1–10.

